# Single-cell profiling of innate and adaptive immune dysregulation in Long COVID

**DOI:** 10.64898/2026.06.04.730206

**Authors:** Sarthak Satpathy, Sean Jordan, Mojtaba Bakhtiari, Jannah Elchommali, Denis Ohlstrom, Siddhartha Mantrala, Ching-Yao Yang, Lahiri Nooka, Tiffany A. Walker, Manoj Bhasin

## Abstract

The COVID-19 pandemic has infected more than 778 million people worldwide. Roughly 7% of these patients progress to Long COVID (LC), suffering from persistent symptoms and cognitive impairment well past the acute stage. As the mechanisms of LC remain elusive, we utilized single-cell profiling (SCP) on 156,478 peripheral blood mononuclear cells (PBMCs) from 20 LC patients and 18 recovered controls (RC) to characterize the disease-associated immune dysregulation. Comparative analysis of LC and RC profiles revealed cellular heterogeneity along with differential abundances across B, T, and myeloid cell compartments. The focused analysis on the B-cell compartment showed that naïve B cells in the LC exhibit elevated IL4R expression and BCR signaling, indicative of sustained antigen exposure and aberrant chronic activation. Concurrently, monocytes adopted heightened interferon signaling and enhanced migratory states, culminating into impaired myeloid differentiation. Furthermore, the T-cell compartment exhibited a functional dichotomy, maintaining sustained quiescence in the central memory compartment while displaying chronic exhaustion within effector memory populations. This dysregulation of effector immunity extended to the NK compartment, where terminally differentiated cells exhibited increased cytotoxicity yet compromised regulatory function, potentially contributing to poor viral clearance. Cellular communication analysis further supports this NK cell dysfunction that is likely driven by galectin and prostaglandin signaling involving monocytes and B cells. We stratified LC patients into mild and severe groups based on symptom and cognitive severity, identifying a distinct immune signature where severe disease is linked to chronic AP-1-mediated inflammation in NK cells and CD14+ monocytes. In contrast, patients with mild symptoms retain functionally competent NK cells with significantly lower exhaustion and apoptosis scores. Collectively, these insights into persistent immune remodeling provide a crucial framework for future biomarker discovery and the development of targeted therapeutic strategies.

**Abstract (Short):** COVID-19 has affected >778 million worldwide, with ∼7% developing Long COVID (LC), characterized by persistent symptoms and cognitive impairment. The mechanisms of LC remain elusive; we utilized single-cell profiling on 156,478 peripheral blood mononuclear cells from LC and recovered controls. Comparative analysis revealed cellular heterogeneity and differential abundance across multiple immune compartments. B-cells exhibited hallmarks of sustained antigen exposure and aberrant activation. Concurrently, monocytes adopted heightened interferon signaling, enhanced migratory states, and impaired differentiation. T-cells exhibited chronic exhaustion within the effector memory populations. Dysregulated effector immunity extended to NK, with increased expression of cytotoxic genes yet compromised regulatory function, potentially contributing to viral clearance. LC patients with severe symptoms showed enhanced AP-1–mediated inflammation in NK cells and CD14+ monocytes, whereas mild cases had fitter NK cells with significantly lower exhaustion and apoptosis. Collectively, these insights provide a framework for biomarker discovery and the development of targeted LC therapeutic strategies.

## Main Text

Globally, the COVID-19 pandemic has resulted in over 778 million confirmed cases as of March 2025^1^. Long COVID or Post-Acute Sequelae of SARS-CoV-2 Infection (PASC) refers to a heterogeneous syndrome characterized by persistent or relapsing symptoms extending beyond the acute phase of COVID-19 illness. Long COVID is estimated to occur in 7% of adults, with potentially less than 10% fully recovering 2 years following infection^2, 3^, underscoring the global public health significance of this condition. A major diagnostic challenge is that Long COVID can emerge even after mild acute illness and in the absence of laboratory-confirmed initial infection.

Long COVID is notable for its clinical diversity, with over 200 reported symptoms affecting multiple organ systems, including respiratory, cardiovascular, neurological, and musculoskeletal systems^4^. Commonly reported features include exercise intolerance (e.g., fatigue, muscle weakness, post-exertional malaise) as well as neurological symptoms such as cognitive impairment (‘brain fog’), dysautonomia, pain syndromes (e.g., myalgia, neuropathic pain, paresthesia), and mood disturbances. While Long COVID can affect individuals regardless of prior health status, certain populations, such as women, individuals from Hispanic and Black communities^5^, and those with severe acute COVID-19 or unvaccinated status ^6^ appear to be at elevated risk.

The pathobiological mechanisms underlying Long COVID remain poorly defined, and effective treatments have yet to be identified. Proposed mechanisms include persistent SARS-CoV-2 reservoirs, immune dysregulation, viral reactivation, and autoimmunity^4^. These diverse possibilities highlight both the complexity of Long COVID and the pressing need for biomarkers to stratify patient risk and guide therapeutic development. Notably, these mechanisms likely involve distinct immune cell types and intercellular signaling networks, which are not adequately captured through bulk tissue profiling. To resolve this complexity at high resolution, we leveraged single-cell RNA sequencing (scRNA-seq), a powerful tool for dissecting the cellular and molecular heterogeneity underlying Long COVID. By profiling gene expression at the single-cell level, scRNA-seq enables the identification of immune cell subsets, transcriptional programs, and signaling pathways that may differentiate patients with persistent symptoms from those who recover fully. Furthermore, this approach can reveal cell type-specific dysregulation in high-risk individuals, potentially uncovering predictive biomarkers and therapeutic targets.

In this study, we performed scRNA-seq on peripheral blood mononuclear cells (PBMCs) from individuals with prior SARS-CoV-2 infection, comparing those with and without persistent long-term symptoms. To understand immune dysregulation, we profiled diverse immune cell subtypes, including B cells, monocytes, and T/NK cells, and examined transcriptional and signaling differences between disease groups. We found evidence of chronic activation and altered antibody production in naive B cells, alongside an elevated migratory and interferon response in monocytes, coupled with depleted myeloid differentiation. Further, the Long COVID related immune dysfunction was characterized by quiescent central memory T cells, exhausted effector memory T cells, and highly cytotoxic but poorly regulated NK cells that demonstrated exhaustion and progression to apoptosis in individuals with high symptom burden. Our results firmly establish Long COVID as driven by persistent, multi-compartment immune dysfunction.

## Results

### Neurocognitive battery and Patient Reported Outcome (PRO) measures reveal significant impairments in individuals with Long COVID

To elucidate the immunological dysregulation underlying Long COVID, the study recruited 38 adults with a history of confirmed COVID-19 (detection of SARS-CoV-2 RNA by nasopharyngeal swab). Participants were categorized into two groups: those reporting new or worsening persistent cognitive symptoms for ≥ 3 months following COVID-19 (Long COVID, N=20) and those without such symptoms (recovered COVID-19, N=16, **Figure 1A).** The study utilized a 38-item Long COVID symptom review and electronic patient-reported outcomes (ePROs) to characterize Long COVID impairments and a feasible telephone-based battery from the National Alzheimer’s Coordinating Center Uniform Data Set to characterize neurocognitive sequelae ^7^.

**Figure 1.**
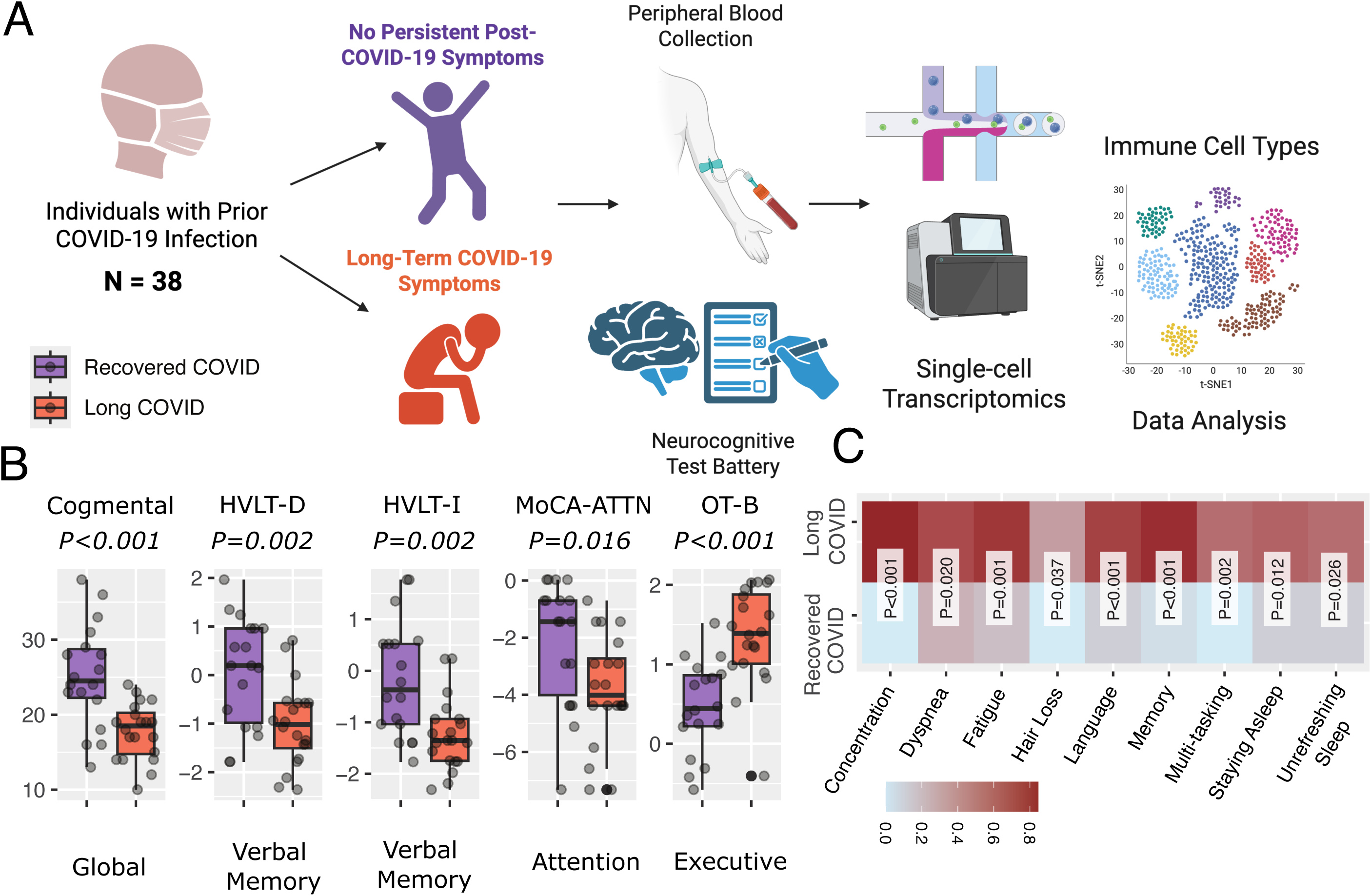
Study design and clinical profiling of individuals with and without Long COVID symptoms. **A. Overview of the study cohort and experimental workflow.** The cohort comprised 38 individuals with a prior history of SARS-CoV-2 infection that were enrolled and stratified based on the presence or absence of persistent post-COVID symptoms. Participants underwent neurocognitive assessment and peripheral blood collection, followed by single-cell RNA sequencing for immune profiling. The figure was prepared using BioRender.com. **B.** Boxplot and scatterplot of telephone-based neurocognitive battery testing scores (*y-axis*) comparing Long (*orange*) and recover COVID (*purple*) across the following cognitive domains (*x-axis*): global function (Cognitive Mental Control, *Cogmental, P<0.001*), verbal memory (Hopkins Verbal Learning Test Immediate, *HVLT-I, P=0.002* and Delayed, *HVLT-D, P=0.002*), attention (Montreal Cognitive Assessment, attention subscale, *MoCA-ATTN, P=0.016*) and attention (Oral Trail Making Part B, *OT-B, P<0.001*). The raw scores were z-score normalized for *HVLT-I, HVLT-D, MoCA-ATTN, and OT-B*. Z-scores are inversely proportional to cognitive function, and the significance of differences in z-scores was assessed by the Wilcoxon rank-sum test. **C.** Heatmap displaying the frequency of dichotomous self-reported symptoms (yes/no) that included concentration difficulties, fatigue, memory loss, language problems, and unrefreshing sleep, among others. Differences between Long COVID and recovered COVID-19 groups were assessed by chi-square and represented by a heat map with red indicating significantly higher symptom frequency.

The two study groups were well-matched on key demographic and clinical characteristics, including age (mean 57.9 vs. 59.8, *P=0.36*), sex (77.8% vs. 90.0% female, *P=0.56*), race (100% African American in both cohorts, *P=1.0*), comorbidity burden (Charlson Comorbidity Index, *P=0.45*), educational level (*P=0.19*), number of prior COVID-19 infections (*P=0.77*), severity of acute illness (*P=0.47*), vaccination status (*P=0.76*), and time since initial infection (*P=0.71*; **Table 1**). Although the Long COVID group showed trends toward greater acute illness severity (50% vs 27.8%, *P=0.47*) and higher rates of Alpha variant infection (40% vs. 16.7%, *P=0.27)* but these findings were not significant.

**Table 1.** Sociodemographic and COVID-related patient characteristics for the study cohort.

|  | Recovered COVID | Long COVID | *P* |
| --- | --- | --- | --- |
|  | (N=18) | (N=20) |  |
| **Gender** |  |  |  |
| Male | 4 (22.2%) | 2 (10.0%) | 0.558 |
| Female | 14 (77.8%) | 18 (90.0%) |  |
| **Age (years)** |  |  |  |
| Mean (SD) | 57.9 (5.24) | 59.8 (6.91) | 0.355 |
| Median [Min, Max] | 57.0 [51.0, 70.0] | 58.5 [50.0, 73.0] |  |
| **Race** |  |  |  |
| African American | 18 (100%) | 20 (100%) | 1 |
| **Charleston Comorbidity Index\*** |  |  |  |
| *Linear scoring, with higher scores demonstrating more pathology |  |  |  |
| Mean (SD) | 3.11 (2.59) | 3.35 (2.62) | 0.779 |
| Median [Min, Max] | 2.00 [1.00, 8.00] | 2.00 [1.00, 9.00] |  |
| **Social Vulnerability Index (SVI)** |  |  |  |
| Low | 0 (0%) | 0 (0%) | 0.454 |
| Medium-Low | 0 (0%) | 1 (5.0%) |  |
| Medium-High | 9 (50.0%) | 7 (35.0%) |  |
| High | 9 (50.0%) | 12 (60.0%) |  |
| **Education** |  |  |  |
| Completed middle school | 0 (0%) | 1 (5.0%) | 0.189 |
| Completed some high school, but did not graduate | 2 (11.1%) | 0 (0%) |  |
| Graduated high school | 5 (27.8%) | 10 (50.0%) |  |
| Completed some college - post-graduate degree | 11 (61.1%) | 9 (45.0%) |  |
| **Presumptive strain\*** |  |  |  |
| *What is the presumptive COVID-19 strain based on date of first infection and predominate circulating strain at that time? |  |  |  |
| Alpha | 3 (16.7%) | 8 (40.0%) | 0.272 |
| Delta | 3 (16.7%) | 3 (15.0%) |  |
| Omicron | 12 (66.7%) | 9 (45.0%) |  |
| **Number of times tested positive for COVID19** |  |  |  |
| 1 | 13 (72.2%) | 15 (75.0%) | 0.773 |
| 2 | 4 (22.2%) | 3 (15.0%) |  |
| 3 | 1 (5.6%) | 2 (10.0%) |  |
| 4 | 0 (0%) | 0 (0%) |  |
| >=5 | 0 (0%) | 0 (0%) |  |

| **Severity of most severe infection** |  |  |  |
| --- | --- | --- | --- |
| No symptoms | 1 (5.6%) | 1 (5.0%) | 0.47 |
| Symptoms without hospitalization | 12 (66.7%) | 9 (45.0%) |  |
| Hospitalized | 4 (22.2%) | 6 (30.0%) |  |
| Intensive Care Unit | 1 (5.6%) | 4 (20.0%) |  |
| Don't Know | 0 (0%) | 0 (0%) |  |
| **Vaccination Status** |  |  |  |
| Mean (SD) | 0.500 (0.514) | 0.450 (0.510) | 0.766 |
| Median [Min, Max] | 0.500 [0, 1.00] | 0 [0, 1.00] |  |
| **Time from initial COVID infection (months)** |  |  |  |
| 0-5 | 0 | 0 |  |
| 6-11 | 3 | 4 |  |
| 12-17 | 3 | 4 |  |
| 18-23 | 1 | 2 |  |
| 24-35 | 8 | 6 |  |
| 36+ | 3 | 4 |  |

Among the various long-term effects of COVID-19, we are particularly interested in neurocognitive sequelae, which remain poorly characterized. As expected, the PROMIS® v2.0 Cognitive Function test showed greater impairment in Long COVID individuals (25.2) compared to the recovered individuals (34.9, *P = 0.0005)*. Neurocognitive testing further revealed significantly reduced performance across multiple cognitive domains in Long COVID patients compared to the recovered group. Specifically, deficits were observed in memory (Hopkins Verbal Learning Test immediate (HVLT-I) and delayed (HVLT-D)) recall, *P=0.0016, P=0.0021*), attention (Montreal Cognitive Assessment attention (MOCA ATTN) subscale score, *P=0.016*), executive function (Oral Trail Making Part B (OT-B) Test, *P= 5.64e-05*), and global cognitive function (Cognitive Mental Control (CogMental), *P=0.0004,* **Figure 1B, Table S1**).

On dichotomous reporting, Long COVID participants experienced higher rates of fatigue, sleep disturbances (e.g., trouble staying asleep, unrefreshing sleep), dyspnea, and hair loss (**Figure 1C, Table S1**). On symptom severity measures, including the modified fatigue impact scale (MFIS), center for epidemiologic studies depression scale (CESD), generalized anxiety disorder scale (GAD-7), and post-traumatic stress disorder checklist (PCL-C), Long COVID participants demonstrated higher impact of fatigue (37.9 vs 21.3, *P=0.0288*), but not depression (20.60 vs 18.8, *P=0.65*), anxiety (16.30 vs 9.83, *P=0.12*), or PTSD (37.9 vs 34.60, *P=0.52,* **Table S1**).

Collectively, these findings define a clinically and demographically well-matched cohort of individuals with and without Long COVID, while uncovering distinct neurocognitive and symptom-based differences. To elucidate the potential contribution of immune dysregulation to the long-term effects of COVID-19, we performed immune profiling using single-cell RNA sequencing.

### Single-cell profiling revealed immune remodeling in Long COVID

Building on the established symptomatic and cognitive differences between participants with and without Long COVID, we next investigated differences in transcriptomic profiles to uncover underlying molecular distinctions between groups. Single-cell profiling captured 83,934 cells from participants with Long COVID and 72,544 cells from the recovered COVID group. Unsupervised clustering and UMAP projection identified seven major immune cell populations, including CD4⁺ T cells (32.74%, *CD3D^+^, CD4^+^)*, CD8⁺ T cells (20.53%, *CD3D^+^, CD8B^+^)*, B cells (11.02%, *CD79A^+^)*, NK cells (9.99%, *CD3D^-^, NKG7^+^)*, CD14⁺ (18.54% *LYZ^+^, CD14^+^)*, CD16⁺ monocytes (6.07% *LYZ^+^, FCGR3A^+^)*, and platelets (<1% *PPBP^+^)* (**Figure 2A-C**). Overall, single-cell transcriptomics successfully captured the major immune cell subsets typically present in peripheral blood mononuclear cells (PBMCs) ^8^.

**Figure 2.**
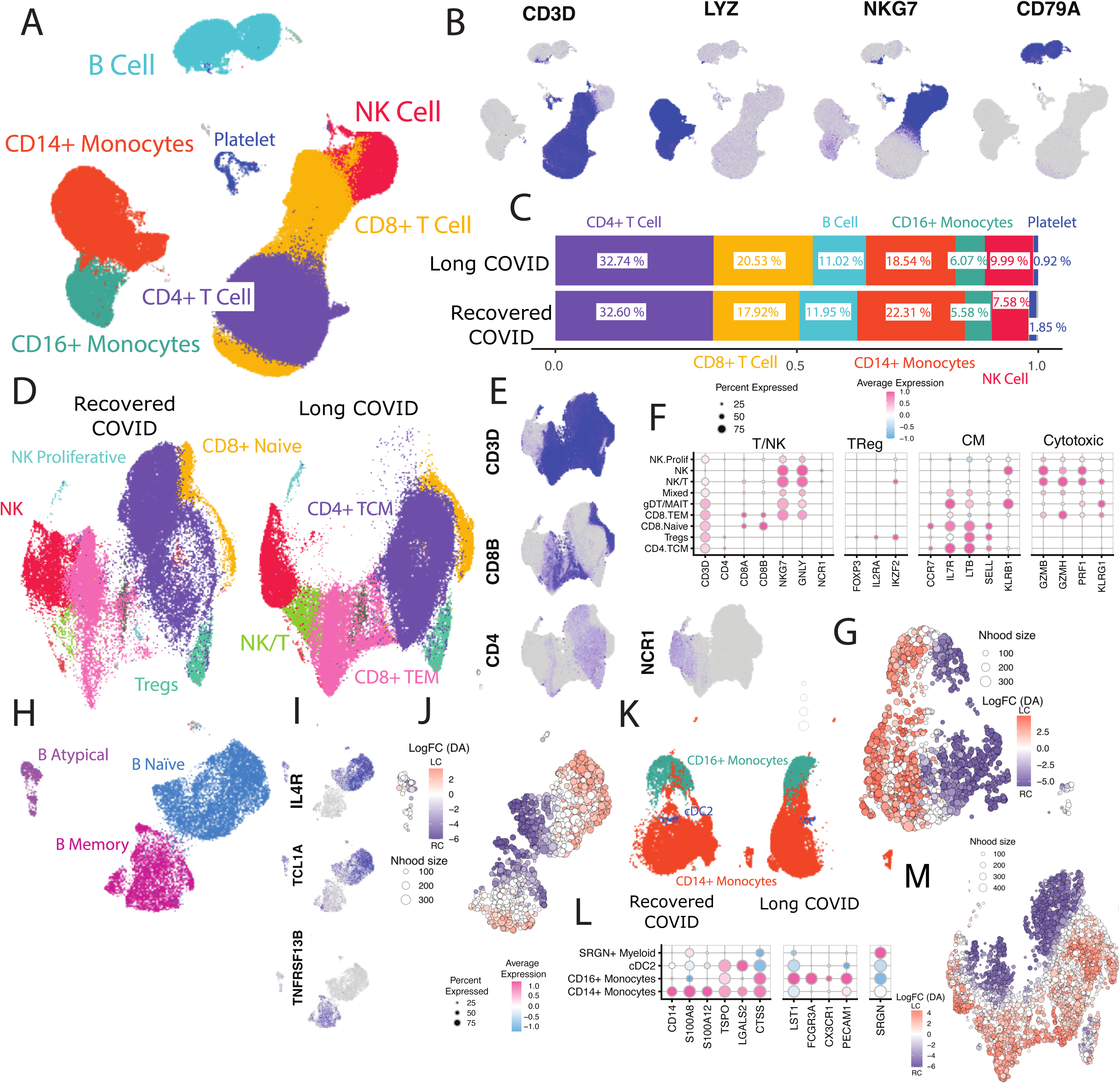
Immune profiling in Long COVID reveals transcriptional remodeling and altered viral repertoire. **A.** Uniform manifold approximation and projection for dimension reduction (UMAP) of all PBMCs from Long COVID and recovered COVID-19 participants, annotated by major immune cell compartments. **B.** Expression of canonical marker genes used to assign immune lineages: *CD3D* (T cells), *LYZ* (monocytes), *NKG7* (NK cells), and *CD79A* (B cells). **C.** Bar plot of proportions of each major immune compartment (CD4^+^ T cell, CD8^+^ T cell, B cell, CD14^+^ Monocytes, CD16^+^ Monocytes, NK cell, Platelet) in Long and recovered COVID groups. **D.** UMAPs of T and NK cells, revealing transcriptionally distinct subsets including CD8⁺ Naive, CD8⁺ TEM, CD4⁺ TCM, Tregs, NK, NK/T hybrids, and NK.Prolif. **E.** Feature plots of CD3D, CD8B, CD4, and NCR1 validating major T and NK subsets. **F.** Dot plots showing average expression of lineage-defining markers across T and NK subtypes. **G.** Neighborhood-level differential abundance map of T/NK compartments; dot size reflects neighborhood size, and color indicates log fold-change (logFC). Higher values of logFC highlight Long COVID enrichment (orange), while negative values highlight recovered COVID-19 enrichment (purple). **H.** B cells’ focused analysis identified B naive, B memory, and B atypical populations. **I.** Expression of TCL1A and IL4R (naive markers) and TNFRSF13B (memory marker) across B cell subtypes. **J.** Neighborhood-level differential abundance analysis of B cells, revealing transcriptionally and condition-specific B cell states. **K.** UMAP of myeloid cells, annotated into CD14⁺ monocytes, CD16⁺ monocytes, and cDC2. **L.** Dot plot of myeloid marker genes confirming cluster identity. **M.** Neighborhood-level differential abundance analysis of myeloid cells, showing differential abundance of transcriptionally distinct neighborhoods in Long- versus recovered-COVID-19 patients.

To gain insight into immune profiles with greater granularity, we independently subclustered the T/NK, B, and myeloid compartments. Using a combination of marker-based analysis and Azimuth reference mapping, we identified immune subtypes of CD4⁺ T, CD8⁺ T, and NK cells with naïve, memory, cytotoxic, and regulatory phenotypes. Identified subsets included CD8⁺ naïve T cells (6.38%, *CD8A^+^, CD8B^+^, CCR7^+^, IL7R^+^)*, CD8⁺ effector memory (TEM) cells (21.13%, *CD8A^+^, CD8B^+^, NKG7^+^, GNLY^+^, GZMB^+^, GZMH^+^),* CD4⁺ central memory (TCM) cells (50.04%*, CD4^+^, CD8B^+^, IL7R^+^, SELL^+^ )*, regulatory T cells ( 3.54%, Tregs, *CD4^+^, FOPXP3^+^, IL2RA^+^, IKZF2^+^ )*, NK cells (12.76%*, CD3D^-^, NKG7^+^, NCR1^+^)*, NK/T hybrids *(3.56*%*, CD3D^+^, NKG7^+^, NCR1^+^, IKZF2^+^ )*, and a proliferative NK population (0.55%*, CD3D^+^, NKG7^+^, MKI67^+^*) (**Figure 2D-F**). The comparative analysis identified distinct subtypes or cell states that were significantly enriched in either Long COVID or recovered COVID-19 patients (FDR < 0.01, **Figure 2G**).

Applying a similar strategy to B cells, we identified three transcriptionally distinct populations: B naïve (61.85%, *CD79A^+^, IL4R^+^, TCL1A^+^),* B memory (31.99%, *CD79A^+^, TNFRSF13B^+^)*, and atypical B cells (4.84%, *CD79A^+^, CD28^+^, CD3D^+^*) (**Figure 2H-I**). Although overall proportions of B cell subsets did not differ significantly between the patient groups, we observed compositional heterogeneity in B-naïve cells between Long COVID and recovered patients (**Figure 2J**).

Examining myeloid cells revealed three main subsets: CD14⁺ monocytes, CD16⁺ monocytes, and conventional dendritic cells (cDC2) (**Figure 2K**). CD14⁺ monocytes (77.15%) expressed *S100A8, S100A12*, and *TSPO*, while CD16⁺ monocytes (21.02%) were marked by *LST1* and *CX3CR1*. cDC2s (1.66%) expressed *LAMP3, PECAM1*. (**Figure 2K**). Differential abundance analysis uncovered transcriptionally distinct CD14⁺ and CD16⁺ monocyte subpopulations exhibiting compositional heterogeneity, highlighting that Long COVID is also associated with innate immune remodeling. (**Figure 2M**).

Cumulatively, these analyses uncovered that while the broad composition of the immune system remained similar between Long COVID and recovered samples, considerable heterogeneity was observed within the respective compartments. To further explore these findings, we next performed a detailed characterization of the different immune compartments.

### Long COVID Naive B-cells Exhibit Increased Co-stimulatory IL-4 Signaling

Building on preliminary evidence of changes in the B cell compartment, we performed a second sub-clustering analysis to further delineate Long COVID-specific changes. (**Fig S1A-C**). The Long COVID group exhibited a broader range of differentially abundant B cell subsets, including enrichment in *IL4+, IGLC1+, and IGLC2+* B naïve as well as *SOX5+* B memory clusters (*P<0.05*, **Figure 3A**). In contrast, the recovered group only showed enrichment in the *ZEB1+* B naïve cluster (*P<0.05*, **Figure 3A**). This suggests more extensive involvement of the B cell compartment in Long COVID-associated immune responses. Using a complementary approach that takes into consideration the cell abundance and fold change, we validated these findings (**Figure 3B)**.

**Figure 3.**
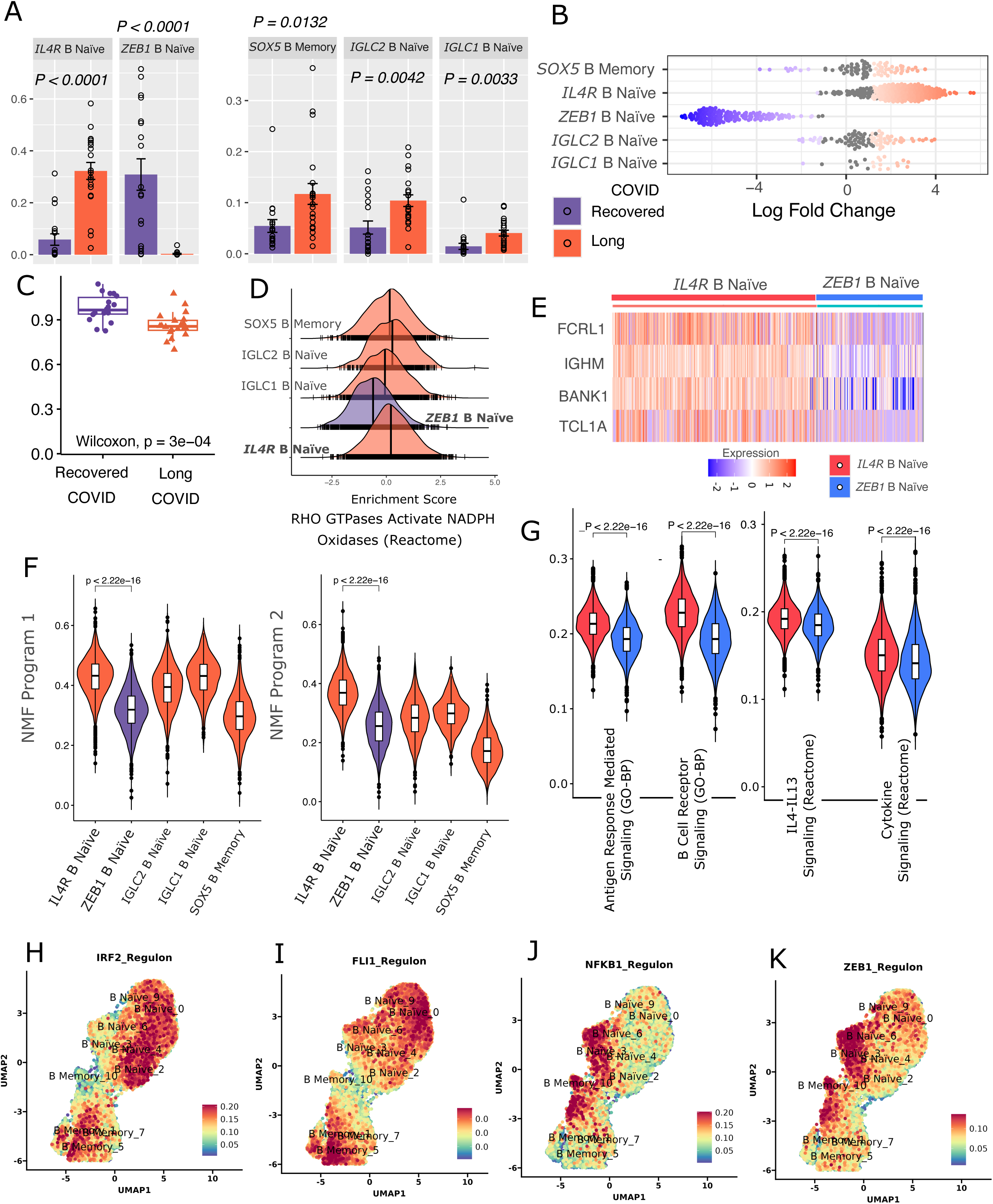
Focused transcriptomic and pathway analysis of the B cell compartment reveals long-term COVID-19-associated alterations in naïve and memory B cells. **A.** Bar plot and scatterplot of the proportion of B-cell subtypes between Long and recovered COVID patients. Bars indicate the mean ± standard error of the proportion for each subtype normalized to the total B cell compartment per patient *(y-axis)*. Statistical significance of the difference was assessed using the Wilcoxon rank-sum test**. B** Beeswarm (scatter) plot from the neighborhood-based differential abundance analysis showing the distribution of log fold changes *(y-axis)* across B cell subtypes *(x-axis)*. Each point represents a small group of transcriptionally similar cells, allowing visualization of subtype-specific abundance differences between recovered and Long COVID patients. **C.** Box plot showing the Shannon diversity index of immunoglobulin (IG) gene usage across B cell subsets. Diversity scores reflect the variability in IG gene expression, with comparisons between recovered and Long COVID groups highlighting potential shifts in clonal diversity. **D.** Ridge (density) plots showing enrichment score (*y-axis*) for RHO GTPase-mediated activation of NADPH oxidases across B cell subtypes. **E.** Heatmap displaying gene expression profiles of differentially expressed genes between *IL4R⁺* naïve B cells (enriched in Long COVID) and *ZEB1⁺* naïve B cells (enriched in recovered COVID). Red indicates high expression, while blue indicates low expression (normalized and scaled). **F.** Violin plots showing enrichment scores for gene modules derived from non-negative matrix factorization (NMF). Two representative modules with distinct expression patterns in *IL4R⁺* and *ZEB1⁺* naïve B cell clusters are shown. Each violin plot illustrates the distribution of module activity from individual cells. **G.** Violin plots showing enrichment scores for four pathways (antigen response mediated, B-cell receptor, IL4-IL13, cytokine signaling) that exhibited strong overlap with gene modules identified by NMF in panel F. **H.-K.** UMAP projections based on gene expression, highlighting enrichment scores for four transcriptional regulons: **H.** IRF2, **I.** FLI1, **J.** NFKB1, and **K.** ZEB1.

Next, to determine whether Long COVID influences immunoglobulin (IG) gene utilization within the B cell compartment, we assessed IG gene usage diversity. Individuals in the recovered COVID-19 group exhibited higher immunoglobulin (Ig) isotype utilization, suggesting a more robust B cell compartment **(Figure 3C**). Pathway analysis revealed Rho GTPase–mediated activation of NADPH oxidase and antigen-driven B cell receptor signaling, culminating in secondary messenger generation within Long COVID–enriched clusters **(Figure 3D, Fig. S1D**). Taken together, these observations along with differential abundance, motivate a comparison between *IL4+* B naïve cluster (enriched in Long COVID) and *ZEB1+*B naïve cluster (enriched in recovered COVID) to elucidate their transcriptional, and functional differences.

*IL4+* B naïve cluster detected higher expression of genes related to B-cell activation or proliferation including *FCRL1,BANK1, TCL1A, IL4R (log2FC >1.5, FDR<0.0001)* and *IGHM* (*log2FC>1*, *FDR<0.0001*) (**Figure 3E, Fig. S1E**). Fc Receptor-Like 6 *(FCRL) 1* plays a role in building the BCR signalosome, influencing B cell responsiveness and antibody production^9, 10, 11^. Along similar lines, B cell scaffold protein with ankyrin repeats (BANK1) participates in B cell signaling, specifically BCR and Toll-like receptor (TLR) pathways, suggesting a role in signaling and may contribute to active defense against latent infection in the Long COVID group^12^. Elevated IL-4 expression in naive cells further supports a role for antigen exposure and subsequent memory B-cell generation in the Long COVID group ^13^. These findings suggest that B cells in Long COVID patients are chronically activated, potentially leading to aberrant immune signaling well after the acute COVID phase has ended.

To identify the key drivers of transcriptomic differences, we examined gene programs active within Long COVID–associated clusters. Two gene programs were highly enriched in the Long COVID naïve cluster (*IL4R+* B naïve) compared to the recovered group (*ZEB1+* B naïve) (**Figure 3F, Fig S1F**). The gene programs showed enrichment for pathways related to antigen receptor–mediated signaling (*FOXP1, IGHM, SKAP1, MEF2C, PDE4D*), IL4-IL13 signaling (*FOS, IL4R, JUNB*) and cytokine signaling (*CAMK2D, FOS, IL4R, JUN, JUNB, LTB*) (**Fig S1G**). Further, module scores for the selected pathways showed higher enrichment in Long COVID dominant IL4R⁺ naïve B cluster compared to its recovered COVID predominant ZEB1⁺ counterpart, indicating distinct pathway activation patterns associated with Long COVID and recovery (*P< 0.0001*) (**Figure 3G)**.

Next, to identify potential transcriptomic mechanisms contributing to differential B cell signaling, we examined the regulatory landscape of naive B cells. We observed distinct patterns of transcription factor activity that differentiate *IL4R+* and *IGLC2+* B naïve (Long COVID high) from *ZEB1+* B naïve (recovered COVID-19 high) clusters (**Fig S1H**). In recovered COVID-19, elevated NFKB1 activity suggests enhanced inflammatory signaling. Additionally, ZEB1 regulon was notably higher enriched in recovered patients, which has been extensively studied in cancer biology but remains poorly understood in the context of chronic viral infection (**Figure 3H-K).**

Collectively, our findings highlight distinct patterns of signaling, transcriptomics, and regulation in naïve B cells from the Long COVID group, potentially indicating a state of chronic activation in response to antigen presentation.

### Classical and non-classical monocytes show higher migratory phenotype in Long COVID

The myeloid compartment, predominantly composed of classical (*CD14⁺*) and non-classical (*CD16⁺*) monocytes that demonstrated distinct clustering patterns between Long and recovered COVID-19 groups **(Figure 2M).** These patterns were largely driven by variations in cell abundance, reflecting group-specific shifts in myeloid cell composition. To better understand the basis of these clustering patterns, we performed focused clustering on the myeloid populations, quantified the relative proportions of each monocyte subtype within the total myeloid cell pool, and annotated the subtypes using canonical marker genes (**Figure 4A, Fig S2A-C,**). This analysis revealed significant shifts in abundance of *CX3CR1^+^ CD16^+^, CX3CR1^-^ CD16^+^*, *IL1B^+^ RETN^Lo^ CD14^+^*, *PDIA3^+^ RETN^Hi^ CD14+* and *DDIT4^+^ PDIA3^+^ CD14^+^*monocytes between individuals with Long COVID and those who had recovered (P<0.05, **Figure 4A**). These changes in cellular composition closely mirrored the group-specific clustering previously observed in B cells, suggesting a convergent mechanism of immune dysregulation driven by altered cell abundance across multiple immune compartments.

**Figure 4.**
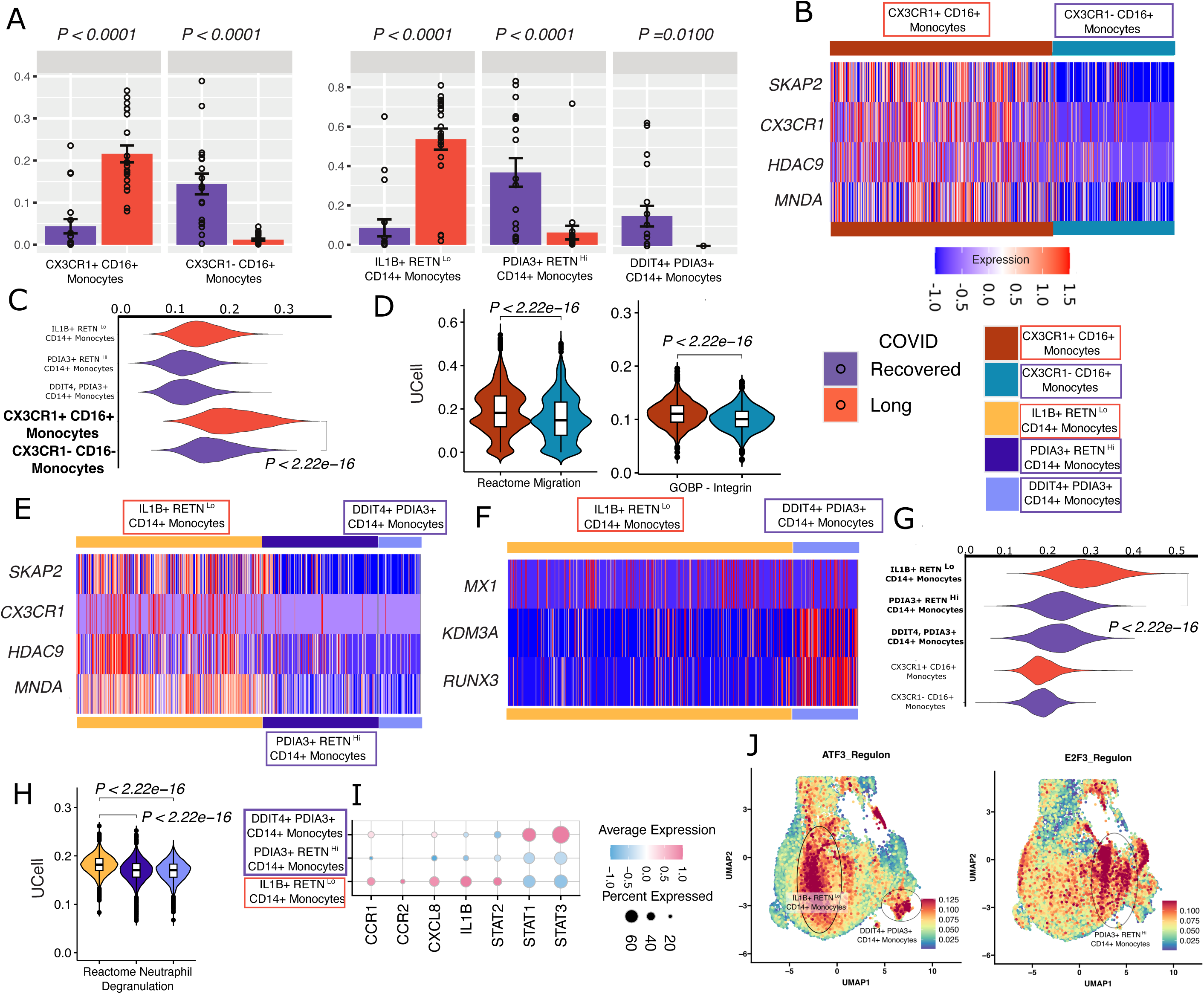
Focused analysis on the myeloid clusters to estimate the effect of Long COVID on classical and non-classical monocytes. **A.** Bar plot and scatterplot of the proportion of monocyte subtypes between Long- and recovered- COVID patients. Bars indicate the mean ± standard error of the proportion for each subtype normalized to the total monocyte compartment per patient *(y-axis)*. Statistical significance assessed using the Wilcoxon rank-sum test**. B.** Heatmap displaying gene expression profiles of differentially expressed genes (*SKAP2, CX3CR1, HDAC9, MNDA*) between *CX3CR1^+^ CD16^+^* monocytes (enriched in Long COVID) and *CX3CR1^-^ CD16^+^* monocytes (enriched in recovered COVID). Red indicates high expression and blue indicates low expression (normalized and scaled). **C.** Violin plots depicting enrichment scores for gene module with distinct scores between *CX3CR1⁺ CD16⁺* and *CX3CR1⁻ CD16⁺* monocyte clusters. **D.** Violin and box plots showing enrichment scores for two pathways – migration (Reactome) and Integrin Signaling (Gene Ontology Biological Process, GOBP) that constitute the NMF gene module from panel C. **E.** Heatmap displaying gene expression profiles of differentially expressed genes (*SKAP2, CX3CR1, HDAC9, MNDA*) in *IL1B^+^ RETN^Lo^ CD14^+^*monocytes (enriched in Long COVID) compared to *PDIA3^+^ RETN^Hi^*and *DDIT4^+^ PDIA3^+^ CD14^+^* monocytes (enriched in recovered COVID). **F.** Heatmap displaying gene expression profiles of differentially expressed genes (*MX1, KDM3A, RUNX3*) in *DDIT4^+^ PDIA3^+^ CD14^+^* monocytes (enriched in recovered COVID) compared to *IL1B^+^ RETN^Lo^ CD14^+^* monocytes (enriched in Long COVID). **G.** Violin plots depicting enrichment scores for gene module with distinct scores in *IL1B^+^ RETN^Lo^ CD14^+^* and *PDIA3^+^ RETN^Hi^* and *DDIT4^+^ PDIA3^+^ CD14^+^*monocytes.**H.** Violin and box plots showing enrichment scores for neutrophil degranulation (Reactome) pathway. **I.** Dot plot displaying the average expression and percent expression of genes associated with inflammatory signaling across *CD14^+^* monocyte subpopulations. **J.** UMAP projections based on gene expression, highlighting enrichment scores for two transcriptional regulons – ATF3 and ETF3.

Among the non-classical monocytes, the *CX3CR1⁺* subset was notably enriched in individuals with Long COVID and exhibited elevated expression of genes such as *SKAP2, CX3CR1, HDAC9* (average log2FC > 1.5, *P<0.001*, **Figure 4A-B**). *SKAP2* drives immune cell migration and adhesion by regulating cytoskeletal dynamics and integrin signaling, which can contribute to inflammatory disease *CX3CR1*, a fractalkine receptor, facilitates both leukocyte migration and adhesion^14^. Further, the elevated expression of HDAC9, a Class IIa histone deacetylase, promotes pro-inflammatory gene expression and is a key regulator in macrophage differentiation^15^. Dysregulation of HDACs more broadly is also linked to autoimmune and chronic inflammatory diseases^16^. While its role in Long COVID remains unclear, its involvement in monocyte trafficking in other chronic conditions suggests it may contribute to inflammatory cell migration in this context as well^17^. We also observe elevated expression of myeloid cell nuclear differentiation antigen (MNDA), which may play a role in controlling pathogen-stimulated type I interferon responses^18^ **(Figure 4B**). Chronic inflammation involves continuous recruitment and activation of immune cells like monocytes in response to a persistent stimulus. Ongoing monocyte influx, combined with impaired terminal differentiation driven by type I interferons, may help sustain the chronic inflammatory response^18^.

We observed a distinct gene program that was differentially enriched in cluster *CX3CR1^+^ CD16^+^* compared to CX3CR1^-^ CD16^+^ non-classical monocytes (**Figure 4C**). Enrichment of the gene program revealed a strong overlap with cell migration (Reactome). Independent analysis of the cell migration pathway confirmed elevated enrichment of migratory signatures in Long COVID samples (**Figure 4D**). Additionally, we observed higher enrichment of integrin signaling pathways, further supporting the role of CX3CR1⁺ non-classical monocytes in trafficking to sites of inflammation (**Figure 4D**). These findings highlight the functional relevance of the identified gene module to immune cell migration and corroborate our differential gene expression results, which also pointed to enhanced migratory and inflammatory potential in Long COVID-associated monocyte subsets.

Moving to classical monocytes, the *IL1B^+^ RETN^Lo^ CD14^+^* cluster was found to be abundant in individuals with Long COVID (**Figure 4A**). This cluster showed upregulation of key genes including *SKAP2, CX3CR1, HDAC9*, and *MNDA*, closely resembling the differential expression of CX3CR1⁺ non-classical monocytes (**Figure 4E**). These genes are associated with enhanced migratory capacity and type I interferon responses, suggesting that both classical and non-classical monocytes may contribute to sustained inflammation and immune cell trafficking in Long COVID. Clusters PDIA3+ RETNHi and DDIT4+ PDIA3+ CD14+ were predominantly enriched in individuals who had recovered from COVID. Comparing cluster DDIT4+ PDIA3+ CD14+ to classical monocytes from Long COVID patients (IL1B+ RETN^Lo^ CD14+) revealed higher expression of genes *MX1, KDM3A,* and *RUNX3*, which are involved in monocyte differentiation (**Figure 4F**). The reduced abundance of DDIT4+ PDIA3+ CD14+ monocytes in Long COVID suggests a lower enrichment of this differentiation-associated phenotype in the disease group.

The IL1B⁺ RETN^Lo^ CD14⁺ classical monocyte cluster, enriched in Long COVID, revealed elevated expression of a gene program related to cytokine signaling (*CD36, CXCL8, IRS2, JUN, NFKBIA, S100A12, CD14,* **Figure 4G, Fig. S2D)**. In addition, genes associated with neutrophil degranulation were also upregulated in the Long COVID abundant classical monocytes (**Figure 4H)**. Collectively, functional enrichment of IL1B⁺ RETN^Lo^ CD14⁺ monocytes indicate heightened innate immune activation and inflammatory potential in Long COVID monocyte populations.

Exploring the interferon response phenotype, we observed distinct interferon signaling activity in both the *IL1B^+^ RETN^Lo^ CD14^+^* and *DDIT4^+^ PDIA3^+^ CD14^+^*monocyte clusters. While both clusters exhibited upregulation of interferon-stimulated genes, the transcriptional drivers differed markedly. The IL1B+ RETNLo CD14+ (Long COVID) cluster was predominantly characterized by elevated expression of genes C*CR1, CCR2, CXCL8, IL1B, STAT2*, whereas the *DDIT4^+^ PDIA3^+^ CD14^+^* (recovered COVID) cluster showed enrichment of genes *STAT1* and *STAT3* (**Figure 4I)**. This divergence highlights the presence of at least two distinct interferon response programs within the monocyte compartment in Long COVID, suggesting functional heterogeneity in antiviral and inflammatory signaling.

Next, we dissected the regulatory landscape underlying the distinct monocyte phenotypes in the Long COVID disease. *IL1B^+^ RETN^Lo^ CD14^+^* (Long COVID) and *DDIT4^+^ PDIA3^+^ CD14^+^* (recovered COVID) monocyte clusters exhibited significantly higher enrichment of the ATF3 regulon (**Figure 4J)**. ATF3 is a stress-responsive transcription factor and a type I IFN–inducible gene that acts as a negative feedback regulator of downstream inflammatory genes ^19^. Supporting this role, ATF3 downregulation in macrophages enhances viral clearance ^20^. Further investigation into ATF3 may clarify mechanisms underlying chronic activation, impaired immune clearance, and neutrophil degranulation (**Figure 4H**). We also found increased enrichment of the FOSB regulon in both clusters, which may reflect a similar inflammatory response^21^ (**Fig S2E)**. However, differences in interferon-stimulated gene expression between these clusters suggest that the interferon response may be regulated differently. In line with earlier findings of reduced monocyte differentiation in Long COVID, we observed lower enrichment of the E2F3 regulon, a transcription factor involved in cell cycle and differentiation (**Figure 4J)**. POU2F2, related to OCT2, was also downregulated, although its role in Long COVID remains unclear (**Fig S2E**. These regulon-level insights support our differential expression results and highlight the transcriptional reprogramming of monocytes in Long COVID.

In summary, we observed an elevated migratory and interferon response phenotype in Long COVID monocytes, along with a depletion of myeloid differentiation.

### Persistent Central Memory T-Cell Quiescence and NK Dysfunction in Long COVID

Having characterized the Long COVID-specific dysregulations within B-cell and myeloid cells, we evaluated alterations in T-cell/NK subset composition, activation states, and regulatory signatures (**Fig S3A-C**). The analysis of the *CD4⁺*central memory T-cell compartment (TCM) revealed five distinct clusters (*LEF1^+^ FHIT^+^ CD4^+^* TCM, *ITGB1^+^ INBP4B^+^ CD4^+^* TCM, *PIK3R1^+^ FTH1^+^ CD4^+^* TCM, *S100A11^+^ S100A4^+^ CD4^+^* TCM, *SNHG7^+^ CIRBP^+^ CD4^+^*TCM), each exhibiting significant differences (*P<0.05*) in abundance between Long COVID and recovered COVID-19 groups (**Figure 5A**).

**Figure 5.**
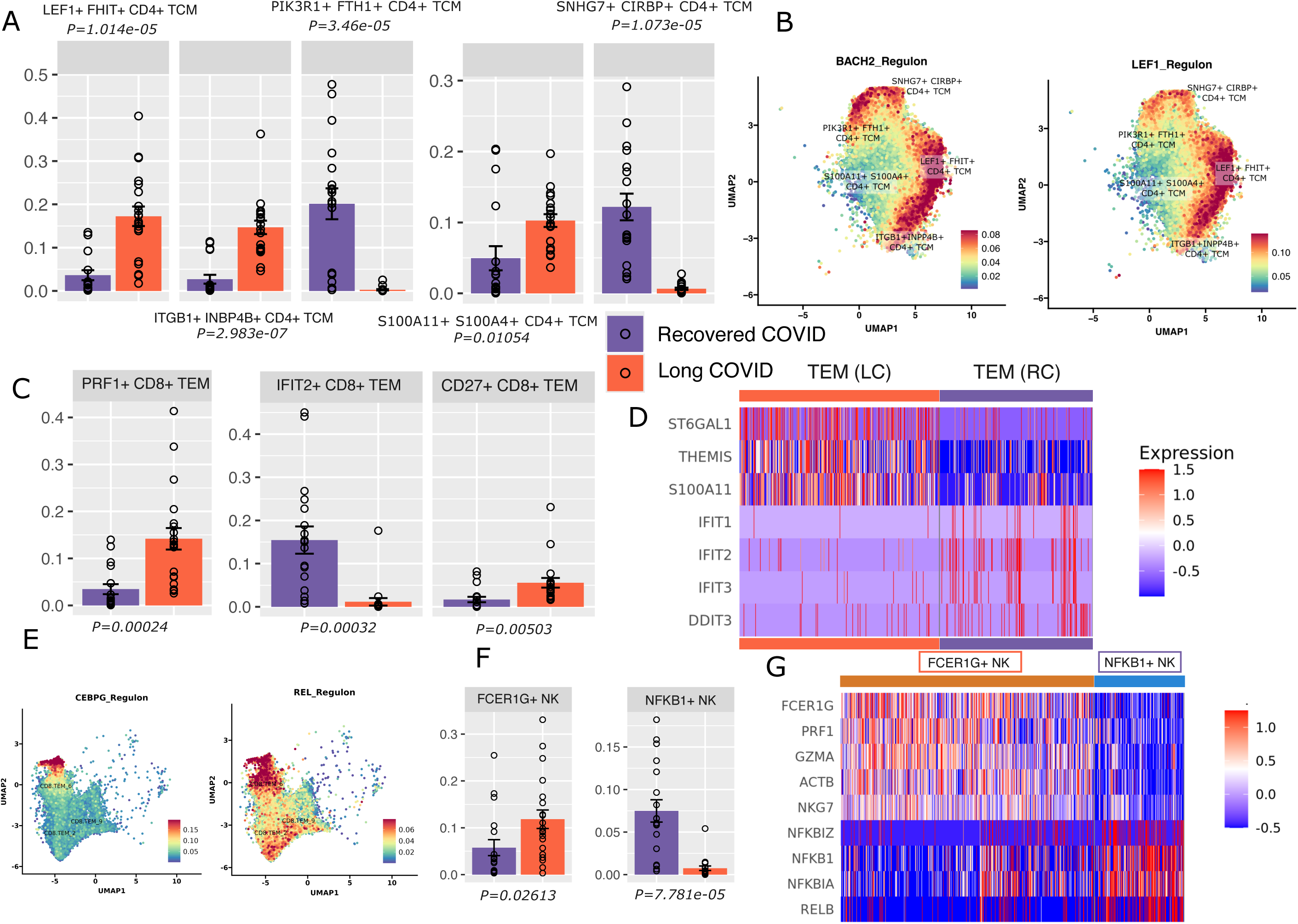
Transcriptional diversity and regulon activity reveal altered T cell states in Long COVID. **A.** Bar plot and scatterplot of the proportion of CD4^+^ T central memory (TCM) cell subtypes between Long and recovered COVID-19 patients. Bars indicate the mean ± standard error of the proportion for each subtype normalized to the total T and NK cell compartment per patient *(y-axis).* Statistical significance assessed using the Wilcoxon rank-sum test. **B.** UMAP projections of CD4^+^ TCM cells based on gene expression, highlighting enrichment scores for two transcriptional regulons: BACH2 and LEF1. **C.** Comparative analysis of the proportion of CD8+ T effector memory (TEM) cell subtypes between Long-and recovered- COVID patients. **D.** Heatmap of differentially expressed genes between Long- and recovered- COVID groups in CD8⁺ TEM cells. Red indicates high expression and blue indicates low expression (normalized and scaled). **E.** UMAP projections of CD8+TEM cells highlighting enrichment scores for two transcriptional regulons: CEBPG and REL. **F.** Comparative analysis of the proportion of NK cell subtypes between Long- and recovered- COVID-19 groups. Bars represent the mean proportion of each subtype normalized to the total T and NK cell compartment per patient. **G.** Heatmap of differentially expressed genes between *FCER1G^+^* and *NFKB1^+^*NK cell subsets. Red indicates high expression, while blue indicates low expression (normalized and scaled).

In the Long COVID group, clusters *LEF1^+^ FHIT^+^ CD4^+^* TCM and *ITGB1+ INBP4B+ CD4+* TCM were more abundant, with significant enrichment of BACH2 and LEF1 regulons related transcriptome profile (**Figure 5B, Fig. S3D**). BACH2 transcription factor is known to suppress the differentiation of central memory T cells into effector memory populations, suggesting a potential block in T-cell maturation ^22^. Further, LEF1 is associated with a cell quiescence program, maintaining T-cell stemness and limiting activation, which may further contribute to the impaired effector differentiation observed in the Long COVID patients ^23^. In contrast, clusters *PIK3R1^+^ FTH1^+^ CD4^+^* TCM and *SNHG7^+^ CIRBP^+^ CD4^+^* TCM were less prevalent in the Long COVID group and showed notable enrichment of the cAMP-responsive element modulator (CREM) transcription factor (**Figure 5B)**. CREM plays a key role in regulating cytokine expression in CD4⁺ T cells, and its enrichment in recovered COVID-19 patients supports the idea of a normal T cell differentiation trajectory in this group^24^. Although DDIT3 was also enriched in the recovered COVID-19 patients that is associated effective antiviral response (**Fig S3D**).

Similar to our findings in the central memory compartment, the T-effector populations (*PRF1^+^ CD8^+^* TEM, *IFIT2^+^ CD8^+^* TEM, *CD27^+^ CD8^+^* TEM) also showed significant differences between Long COVID and recovered COVID groups. **(Figure 5C).** In the Long COVID group, TEM cells showed elevated expression of *ST6GAL1*, *THEMIS*, and *S100A11* **(Figure 5D)**. ST6 Gal I has been shown to promote rapid IL-2Rα expression and early proliferation in terminal effector CD8⁺ T, suggesting a hyperactivated phenotype ^25^. THEMIS, on the other hand, is known to suppress effector function during acute viral infections, indicating a potential regulatory or dysfunctional state^26^. *S100A11*, a calcium-binding protein, may be associated with an exhaustive phenotype in Long COVID T cells^27^. In contrast, the recovered COVID-19 group exhibited higher expression of interferon-stimulated genes, including *IFIT1, IFIT2,* and *IFIT3* **(Figure 5D)**. Additionally, DDIT3 was upregulated, which may reflect a more effective antiviral response and resolution of inflammation in the recovered COVID-19 group. These transcriptional differences highlight distinct effector T-cell programs in Long COVID, characterized by altered activation, regulation, and exhaustion based on overexpression of S100A11 gene.

Furthermore, we observed elevated activity of transcription factors related to CEBPG and REL in the recovered COVID-19 group across the effector T cell compartment **(Figure 5E**). These transcriptional regulators are key modulators of both innate and adaptive immunity and play essential roles in maintaining immune homeostasis. Their increased expression in recovered individuals suggests a coordinated regulatory mechanism that supports effective immune resolution and protects against chronic inflammation. This complements our earlier findings of enhanced interferon signaling and effector function in the recovered group, highlighting a more robust and regulated immune response compared to the dysregulated profiles observed in Long COVID **(Figure 5E**).

Beyond the T-cell compartment, our investigation into NK cell populations revealed two differentially abundant subsets with distinct transcriptional profiles in Long COVID and recovered COVID groups **(Figure 5F).** The *FCER1G+* NK cluster, which was more prevalent in Long COVID, exhibited elevated expression of cytotoxic molecules, including *FCER1G, PRF1, GZMA, ACTB, PNC1,* and *NKG7* **(Figure 5G)**. This expression profile is indicative of a mature and terminally differentiated NK cell population, suggesting sustained cytotoxic activity in Long COVID^28^. In contrast, the *NFKB1⁺* NK cluster, predominant in recovered COVID-19 individuals, expressed both inflammatory and anti-inflammatory genes, including *NFKB1, NFKBIA, NFKBIZ, and RELB* **(Figure 5G)**. These genes, linked to NF-κB signaling, are known to promote immune cell recruitment such as T cells ^28, 29^. This expression pattern suggests that NK cells in recovered individuals maintain a homeostatic yet alert state, balancing activation with regulation to facilitate coordinated immune resolution.

In summary, our results reveal distinct immune signatures in Long COVID compared to recovered COVID. Long COVID is marked by sustained T-cell quiescence in the central memory compartment, chronic exhaustion in effector memory T cells, and terminally differentiated NK cells with heightened cytotoxicity but limited regulatory function. In contrast, recovered COVID shows active transition from central to effector memory T cells, effective effector responses, and NK cells enriched for inflammatory and regulatory genes, supporting immune resolution and viral clearance.

### Galectin and Prostaglandin Signaling Drives immune dysregulation and NK Impairment

To further understand the interplay of immune cell types driving dysregulation in Long COVID, we performed a cell-cell communication analysis focusing on key lymphocyte and monocyte populations that were significantly associated with either recovered or Long COVID. A notable finding was galectin-mediated signaling originating from monocytes and targeting NK cells, observed exclusively in the Long COVID patients (**Figure 6A**). This pathway likely promotes NK cell dysfunction, potentially through the expression of the ligand LGALS9 on monocytes and its corresponding receptor P4HB on *FCER1G⁺* NK cells (**Figure 6B**) ^30^. P4HB receptor, which has been shown previously to upregulate in the setting of viral infections, was expressed only in *FCER1G⁺*NK cells from Long COVID samples, suggesting a Long COVID-specific ligand-receptor interaction that may drive terminal NK cell differentiation and limit their regulatory capacity (**Figure 6B**) ^31^.

**Figure 6.**
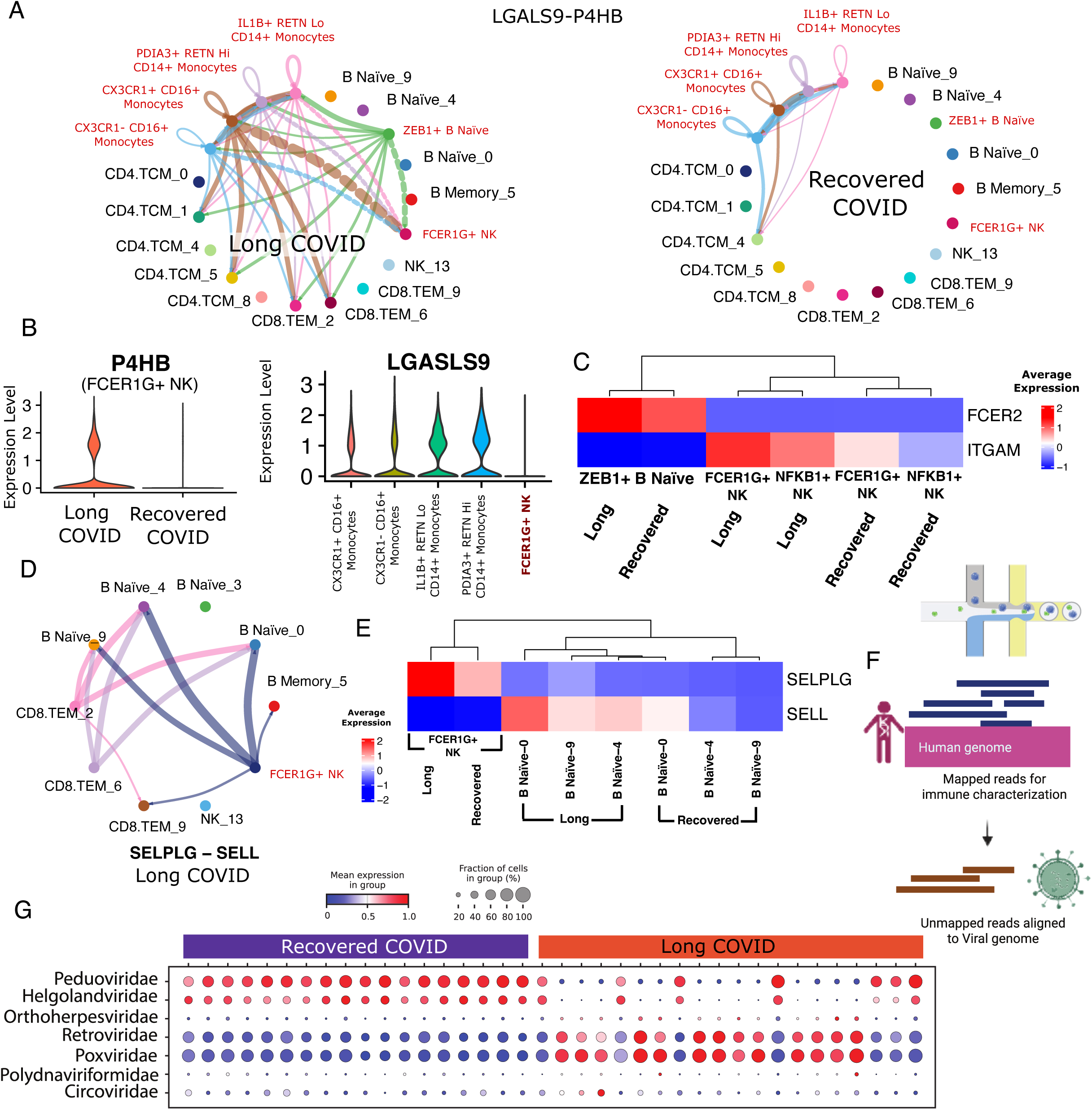
Altered intercellular communication networks across immune compartments in Long COVID. **A.** Circle plot illustrating cell-cell communication via the galectin signaling pathway, mediated by the ligand LGALS9 and receptor P4HB. The width of each edge reflects the inferred strength of interaction between cell types. This visualization highlights altered intercellular signaling dynamics in Long COVID (left) compared to recovered COVID-19 (right). **B.** Left: Violin plots showing expression of the receptor P4HB on *FCER1G⁺*NK cells in Long- and recovered- COVID-19 groups, highlighting disease-associated differences in receptor expression. Right: Violin plots comparing expression of the ligand LGALS9, which is exclusively expressed by monocytes, across CD14⁺ monocytes, CD16⁺ monocytes, and *FCER1G⁺* NK cells. These plots illustrate the cell-type–specific distribution of ligand-receptor components involved in galectin signaling. **C.** Heatmap of pseudo bulked expression of the ligand FCER2 and receptor ITGAM, components of the CD23 signaling pathway, across *ZEB1^+^* B naïve cells, *FCER1G^+^* NK cells, and *NFKB1^+^* NK cells in Long- and recovered- COVID groups. **D.** Circle plot illustrating cell-cell communication via the SELPLG signaling pathway, mediated by the ligand SELPLG and receptor SELL in Long COVID patients. **E.** Heatmap of pseudobulked expression of the ligand SELPLG and receptor SELL, components of the SELPLG signaling pathway, across *FCER1G^+^*NK cells, *IL4R^+^* B, *IGLC1^+^* B, *IGLC2^+^*B naive cells in Long COVID and recovered COVID groups. **F.** Schematic of viral repertoire estimation pipeline: unmapped reads from scRNA-seq were re-aligned to a viral genome database to assess viral content. The figure was prepared using BioRender.com. **G.** Dot plot showing abundance and expression of viral families across samples. Retroviridae and Poxviridae were enriched in Long COVID (*log2FC > 1.25, P<0.0001*).

Furthermore, B cells in Long COVID displayed increased FCER2 expression, which may facilitate their interaction with NK cells via the ITGAM receptor, potentially modulating NK cell function again. Although *ZEB1⁺* B cells showed only a slight increase in FCER2 ligand expression, ITGAM receptor levels were markedly higher in both *FCER1G⁺* and *NFKB1⁺* NK cells in Long COVID patients. This enhanced ligand-receptor interaction is likely to promote NK cell cytotoxicity, contributing to chronic inflammation without effective immune clearance ^32^(**Figure 6C**).

SELPLG-mediated signaling from *FCER1G⁺* NK cells to naïve B cells emerged as another pathway exclusively associated with Long COVID (**Figure 6D**). This interaction was supported by elevated expression of the SELPLG ligand in NK cells and increased expression of its receptor, SELL, in B cells (**Figure 6E**). Although the functional impact of this pathway remains unclear, it may represent a novel axis of immune crosstalk contributing to altered B cell regulation in Long COVID, warranting further investigation ^33, 34^. Further secondary validation through metagenomic analysis revealed consistent enrichment of specific viral families in Long COVID samples, including Retroviridae and Poxviridae **(Figure 6F-G)**, both at abundance and expression levels. These findings indicate that altered virome content may contribute to the dysregulated Long COVID immune landscape.

Collectively, cellular communication analysis highlighted a reprogrammed immune signaling network in Long COVID, defined by selective crosstalk among monocytes, B cells, and NK cells. The emergence of disease-specific ligand-receptor interactions suggests a unique immunoregulatory axis that may contribute to persistent inflammation and impaired immune resolution.

### Innate Immune Dysregulation Underlies Long COVID Symptom Severity

To identify immune features associated with Long COVID symptom severity, we performed a weighted correlation analysis between cell type proportions and patient-reported symptom burden. This revealed a significant inverse correlation between the proportion of NK cells and symptom load, and a positive correlation between CD14+ monocytes and symptom load (**Figure 7A**). To understand whether NK cell abundance reflected functional differences in the cohort, we next assessed exhaustion and apoptosis scores in this population. NK cells from patients with more severe Long COVID exhibited significantly higher exhaustion and apoptosis scores compared to those from less severe cases (**Figure 7B**). In contrast, there were no significant differences in the distribution of cell cycle phases between the two groups (**Figure 7C**), suggesting that NK cell depletion in severe Long COVID is likely driven by terminal dysfunction and cell death rather than impaired proliferation. Supporting this interpretation, NK cells in less severe cases showed increased enrichment of the electron transport chain and oxidative phosphorylation (OXPHOS) pathways (**Figure 7D**), consistent with a metabolically active and functionally competent immune phenotype. Gene regulatory network analysis further revealed upregulation of JUND, FOS, and IRF1 regulons in NK cells from severe Long COVID patients, indicative of an inflammatory and stress-responsive transcriptional program. In contrast, CEBPD, a transcription factor known to promote metabolic fitness in NK cells ^35^, was selectively upregulated in NK cells from less severe patients (**Figure 7E**). Together, these findings suggest that preserved NK cells in less severe Long COVID patients exhibit features of metabolic and transcriptional resilience, while NK cells in more severe Long COVID are transcriptionally rewired into an inflammatory and dysfunctional state.

**Figure 7.**
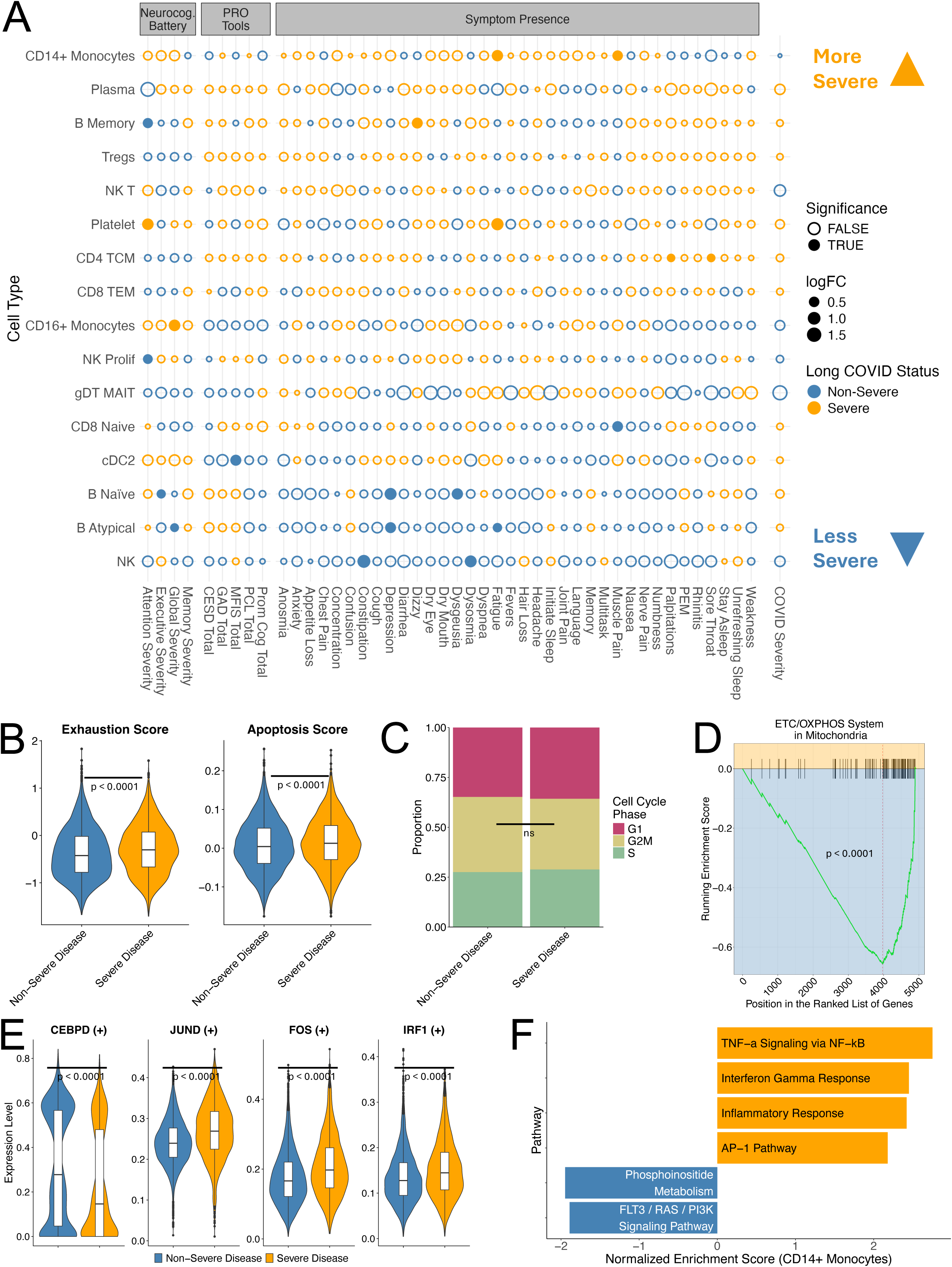
NK cell exhaustion and innate immune dysfunction stratify long COVID symptom severity. **A.** Correlation heatmap showing associations between cell type proportions and patient-reported symptom scores across individuals with long COVID. Dot size reflects the magnitude of effect, and open circles indicate statistically significant associations (FDR < 0.05). NK cell abundance negatively correlates with symptom burden, while *CD14*⁺ monocytes show a positive correlation. **B.** Violin plots showing significantly higher exhaustion and apoptosis scores in NK cells from patients with severe long COVID, suggesting terminal dysfunction. **C.** Barplot showing no significant difference in the distribution of NK cell cycle phases between disease severity groups, indicating that the loss of NK cells in severe cases is not due to impaired proliferation. **D.** GSEA plot demonstrating enrichment of electron transport chain and oxidative phosphorylation (OXPHOS) pathways in NK cells from patients with less severe symptoms, consistent with metabolic resilience. **E.** Violin plots showing increased expression of the regulatory transcription factor CEBPD in NK cells from less severe patients, while AP-1-related stress-responsive factors JUND and FOS, and the interferon-associated IRF1, are upregulated in severe long COVID. **F.** GSEA of CD14⁺ monocytes reveal that pro-inflammatory pathways (e.g., TNF-α signaling, IFN-γ response, AP-1 signaling) are enriched in severe long COVID, while pathways related to immune regulation and survival (e.g., FLT3 / RAS / PI3K signaling and phosphoinositide metabolism) are upregulated in less severe cases.

We observed a similar bifurcation in CD14+ monocytes, where gene set enrichment analysis revealed that TNF-α signaling via NF-κB, interferon-γ response, and AP-1-driven inflammatory pathways were upregulated in monocytes from severe Long COVID patients. In contrast, monocytes from less severe patients showed significant enrichment for phosphoinositide metabolism and FLT3 / RAS / PI3K signaling pathways (**Figure 7F**), which are involved in inflammation resolution and immune regulation^36^. These findings mirror the dichotomy observed in NK cells and suggest that Long COVID severity is associated with coordinated dysregulation of multiple innate immune cell types.

Taken together, these results support a model in which chronic AP-1-mediated inflammation, sustained by both NK cells and CD14+ monocytes, contributes to the pathogenesis of severe Long COVID. Therapeutic strategies aimed at modulating AP-1 activity, restoring immune metabolic resilience, or disrupting pro-inflammatory crosstalk between innate immune populations may provide novel avenues for intervention.

## Discussion

Given Long COVID heterogeneity and the complexity of symptom-based primary endpoints in current clinical trials, there are ongoing efforts to characterize inciting factors and pathobiological mechanisms driving Long COVID to identify intervenable targets for drug repurposing and development. Uniquely, we provide a detailed landscape of gene programs and gene regulatory networks guiding innate and adaptive immune dysregulation in a well-characterized Long COVID population with diverse symptom profiles and a high burden of cognitive impairment. We observed differential abundance within immune cell subsets, with distinct populations of CD8+ naïve T cells, CD8+ TEM, CD4+ TCM, proliferative NK cells, naïve and memory B cells, and CD14+ and CD16+ monocytes. Leveraging single cell profiling, we describe an inflammatory stress response supporting the leading theory of persistent antigen exposure^37^, as evidenced by gene regulation and expression driving 1.) T cell exhaustion and dysregulated differentiation, 2.) enhanced NK cell cytotoxicity and apoptosis 3.) BCR-mediated proliferative naïve B cells, and 4.) migratory monocytes with increased interferon and cytokine signaling and impaired differentiation.

Long COVID was marked by increased proliferation of naïve and memory B cells, consistent with previous findings of SARS-CoV-2- specific naïve B cell activation with elevated antibody production against SARS-CoV-2 spike and nucleocapsid antigen, all supporting an ongoing response to a persistent viral reservoir ^38, 39, 40, 41, 42^. This is likely to occur, in part, through Long COVID-specific differential expression of *FCRL1,* which peaks in naïve and memory B cells in response to BCR activation, leading to BCR signalosome formation, B cell proliferation, and amplified humoral responses ^9^. Similarly, differential expression of BANK1 in response to BCR engagement enhances B-cell signaling, and multiple gene programs enriched here are linked to cytokine and antigen-receptor signaling^12^. We demonstrate increased IL-4 expression and IL 4-IL 13 signaling axes in naïve B cells. This pathway has been implicated as a driver of severe acute COVID-19, and neutralization of this pathway with IL4-IL13 specific monoclonal antibody has reduced disease severity ^43^. IL4-IL13 signaling aligns with evidence of ongoing antigen exposure and likely contributes to memory B-cell generation and stimulation of B cell differentiation with subsequent antibody production^44^. PSGL-1 mediated signaling between NK cells and naive B cells observed in Long COVID participants may contribute to B cell activation and cell trafficking to affected sites ^34^. IRF2 enrichment in Long COVID further indicates a proliferative and antibody-producing phenotype and may correlate with interferon-mediated pathways, as demonstrated in our monocyte population and as shown previously ^45^. ^37^ Notably, continued activation of naïve B cells in this cohort occurs at a much later Long COVID duration than in previous studies, highlighting the durable and protracted nature of antigen-mediated immune responses following SARS-CoV-2 infection. This contributes to the growing body of evidence highlighting the importance of identifying culprit viral-specific antigens, locating the reservoir, and developing strategies for eradication.

We observed differential abundance and disease-specific gene expression within CD14+ and CD16+ monocytes that supports antigen-dependent migration, evidenced by enrichment of *SKAP2, CX3CR1, HDAC9,* and further corroborated by NMF gene programs for Reactome migratory and integrin pathways in CD16+ monocytes ^14, 15, 16^. These findings align with previous studies showing increased expression of CXCR6 and PSGL-1 in Long COVID, proposed to promote monocyte recruitment in response to ongoing inflammation in organ tissues, such as the lung ^46^. We demonstrate that CD16+ and CD14+ monocyte clusters express *MNDA*, a gene associated with pathogen-stimulated type I interferon responses often seen in response to viral pathogens ^17, 18^. The distinct interferon signaling patterns we observed support previous reports of IFNβ upregulation in Long COVID ^38^. Inflammatory signatures are seen through cytokine signaling and toll-like receptor cascades in CD14+ monocytes that likely trigger enrichment of the ATF3 regulon, which regulates stress responses and may contribute to ineffective viral clearance ^19, 20^. In addition, there was also a lower abundance of *MX1, KDM3A,* and *RUNX3* expression and lower enrichment of the E2F3 transcription factor, indicating decreased monocyte differentiation in Long COVID ^47^. Together, these findings support a persistent response to a distal, potentially tissue-based antigen reservoir, in keeping with current mounting evidence ^37^.

The most significant alteration to T cell subtypes in Long COVID was observed in T memory cells, with gene expression suggesting quiescent CD4+ TCM cells that fail to differentiate into effector memory cells and populations of CD8+TEM cells undergoing terminal differentiation with impaired effector function and exhaustion phenotypes ^22, 23, 24^. This process is associated with CD4+ and CD8+ activation and exhaustion, evidenced previously by PD1 and TIM3 expression ^41, 48^, and here by THEMIS and S100a11 expression ^26, 27^. In contrast, the recovered COVID-19 group displayed enrichment of transcription factors related to CREM, CEBPG and REL that support normal differentiation and homeostatic functions in this population and may represent more effective immune regulation ^24, 49^. Similarly, NK cells have been shown to have an overactive cytotoxic phenotype in Long COVID ^50^. To further this understanding, we describe terminally differentiated NK cells with impaired cytotoxic profiles in contrast to recovered COVID-19 NK cells, which expressed a diverse pattern of pro-inflammatory genes that may effectively aid in communication and recruitment of other immune cells, such as T cells. Although previous studies did not isolate exhaustion patterns in this population ^50^, we describe expression of exhaustion markers, metabolic effector uncoupling, and increased progression to apoptosis in the severe Long COVID group. This phenotype may lead to ineffective viral clearance and may be driven by galectin- and ITGAM-mediated cell communication was observed from monocytes and naive B cells to NK cells ^32^. Collectively, the dysregulated and exhausted patterns observed in Long COVID participants provide further support for persistent antigen exposure, as these responses can be seen during chronic infection ^51, 52, 53^.

Due to variable symptom presentation in Long COVID, there has been limited success in defining distinct phenotypes and linking these manifestations with associated pathobiological mechanisms. While Long COVID is broadly associated with persistent NK cell dysfunction compared to recovered individuals, our intra-cohort analysis revealed the opposite trend: higher NK cell abundance correlated with less symptom burden among Long COVID patients. This reinforces the idea that NK cell presence alone is not inherently protective or pathogenic but must be interpreted in the context of their functional and metabolic state. This aligns with our observation that NK cells in less symptomatic patients exhibit lower exhaustion and apoptosis scores, as well as greater metabolic activity, suggesting they may be functionally competent and metabolically resilient. This supports a model in which protective NK subsets resist terminal exhaustion and help sustain immune equilibrium to retain effective antiviral function. In contrast, patients with more severe symptoms appear to harbor NK cells that are transcriptionally rewired into a dysfunctional and inflammatory state. This layered immune dysregulation parallels phenomena observed in chronic viral infections such as Hepatitis B and C, where prolonged antigenic stimulation drives NK cell exhaustion and contributes to tissue pathology, while regulatory or metabolically resilient NK cells help mitigate damage ^54^. Therapeutically, this distinction is critical. Strategies that broadly suppress NK activity could disrupt protective subsets and exacerbate immune dysfunction. Instead, targeted interventions that restore metabolic homeostasis, reduce chronic AP-1-driven inflammatory signaling, or reprogram exhausted NK cells may offer more effective avenues for alleviating Long COVID severity.

Although persistent SARS-CoV-2 viral reservoir and reactivation of previous viral infections are leading hypotheses of etiologic contributors to immune dysregulation in Long COVID, it remains unclear which viral antigens are triggering this process and in which individuals. Persistent antigen and elevated antibody titers have been detected for both SARS-CoV-2 and herpesviruses, particularly Epstein-Barr Virus (EBV) ^51, 52, 53^. However, detection of SARS-CoV-2 RNA or herpesvirus DNA from peripheral blood has been limited across studies, and it is equivocal whether this is an effective way of measuring viral activity if confined to a tissue reservoir. Similarly, investigation of persistent viral elements from peripheral blood in our study revealed no significant difference in Coronoviridae or Herpesviridae. However, we demonstrate differential abundance of terminally differentiated TEM CD8+ cells, which has been previously significantly correlated with EBV antibody reactivity on REAP ^40^. Further investigation is needed to identify if these viruses are contributors to the immune dysregulation seen in this study and whether peripheral blood sampling limits detection of pathogenic viral particles sequestered to tissue reservoirs.

Uniquely, we evaluated immune dysregulation in a population of African Americans with high social vulnerability, a demographic underrepresented in Long COVID pathobiological research ^40, 41^. COVID-19 has disproportionally affected African American communities with significantly higher morbidity and mortality rates, and recent evidence suggests that social risk factors and systemic barriers increase the risk of Long COVID development and persistence ^55, 56^. Long COVID profiles may vary by race and ethnicity, indicating potential variation in the underlying drivers of disease versus variation in the symptom manifestation of those drivers ^57^. Our overarching findings are consistent with the latter and support current evidence of immune activation and fatiguability in response to persistent antigen exposure.

Similarly, neurocognitive phenotypes are less commonly studied compared to persistent pulmonary symptoms; however, we identified similar immune profiles to those reported previously, with expanded detail of the transcriptomic landscape to promote biomarker discovery. Current evidence suggests these immune mechanisms likely contribute to microglial activation following blood-brain barrier disruption, triggering cognitive impairment ^58, 59^, but further investigation is needed to comprehensively characterize this process. Notably, cognitive impairment did not appear to be driven by mental health disorders in our population, further supporting an independent pathological process as has been proposed previously ^60^.

There are several limitations of this study. First, convenience sampling may lead to bias in our cohort; however, we did not observe significant differences in sociodemographic and clinical characteristics between Long COVID and recovered COVID-19 participants. The time range between the incidence of COVID-19 and enrollment in the study varied. Although there is some evidence that Long COVID mechanisms may evolve throughout the course of the condition ^61^, we observed similar mechanisms across the cohort despite varying Long COVID durations. We identify gene expression patterns of NK and CD8+ T cell exhaustion; however, these findings should be validated and formally assessed through functional assays. We observe mechanisms consistent with antigen-mediated inflammation, but the inciting antigen remains unclear and should be investigated further through T cell receptor sequencing. Peripheral blood sampling allowed characterization of these immune responses, including evidence of cell migration to affected tissues, but further investigation of host tissues is needed to identify viral reservoirs and characterize focal immune responses.

In summary, we leveraged single-cell profiling to comprehensively characterize immune signatures corresponding with Long COVID symptom persistence compared with well-matched COVID-19 recovery in African Americans. We demonstrate both adaptive responses and immune dysfunction consistent with protracted viral antigen exposure. We detail several immune processes that may aid in biomarker generation or serve as modifiable targets for drug development.

## Methods

### Study population

We recruited African American adults aged ≥50 years from Grady Memorial Hospital in Atlanta, GA between December 2023–December 2024 with a history of confirmed SARS-CoV-2 test who reported either a.) new or worsening symptoms lasting ≥3 months (Long COVID) or b.) no persistent symptoms (recovered COVID-19 following SARS-CoV-2 infection. All blood samples were collected at the time of enrollment. The time interval between participants’ first COVID infection to blood collection for the study ranged from 7 months to 53 months (**Table 1**). No participant was sampled at multiple time points. All data points represent unique participants. Ethics approval was obtained from Emory University IRB (IRB5939) prior to study initiation, and written consent was obtained as indicated.

### Data Collection and Case Definitions

Sociodemographic (age, sex, self-reported race, SVI), comorbidities, and acute COVID-19 data (disease onset, severity, and vaccination history) were collected by survey and confirmed by electronic health record review. As outpatient oxygen saturation stratification was not consistently available for mild acute COVID-19 cases, we defined mild cases as acute COVID-19 not requiring inpatient hospitalization within 14 days of symptom onset and severe cases as requiring inpatient hospitalization or intensive care unit (ICU) admission for COVID-19 complications within 14 days of symptom onset. Presumptive strain was defined by the predominant circulating strain at the time of the participant’s SARS-CoV-2 infection. Symptoms experienced within the last 7 days were collected with dichotomous scoring (presence/absence). To estimate the severity of prevalent Long COVID symptoms, participants completed validated patient-reported outcome (PRO) tools including the PROMIS Cognitive Function 8A ^62^, Modified Fatigue Impact Score (MFIS) ^63^, Center for Epidemiologic Studies Depression Scale (CESD) ^64^, and Generalized Anxiety Disorder (GAD-7) ^65^, and PTSD Checklist (PCL-C) scales. Study data were collected and managed using REDCap electronic data capture tools hosted at Emory University ^66^.

Balancing burden to a traditionally socioeconomically disadvantaged population, all participants underwent a comprehensive telephone-based neurocognitive battery that evaluated the following cognitive domains: global function (Montreal Cognitive Assessment (MOCA) Blind), executive function (Number span backward, Oral Trail Making Part B), memory (Craft Story immediate and delayed, Hopkins Verbal Learning Test immediate (HVLT-I) and delayed (HVLT-D)), and attention (Number span forward and MOCA Blind Attention subscale) ^7, 67, 68, 69, 70^. Raw scores on the cognitive tests were converted to z scores based on the means and SDs of demographically comparable normative samples.

### PBMC isolation and single-cell preparations

PBMCs were isolated using the standard Ficoll-Paque density-gradient method according to the manufacturer’s instructions (Cytiva). Briefly, blood diluted in phosphate buffer saline (PBS) (1:1) was gently layered onto Ficoll-Paque PLUS (Cytiva, 17144002) and spun at 500g for 30 minutes at 21°C. The top layer (plasma) was removed and discarded. The layer containing the mononuclear cells was then carefully removed and diluted with 3x volume of PBS, mixed well, and spun at 500g, for 10 minutes at 21°C. The pellet was resuspended in PBS and washed again by spinning for 10 minutes at 500g and 21°C. The PBMC pellet was then resuspended in 1 ml recovery cell culture freezing media (Fisher Scientific, 12648010) at a concentration of <10X106 cells/ml. Frozen PBMC samples were thawed and washed with wash buffer (PBS containing 1% BSA) to prepare viable single-cell suspensions

### Single-cell assays and sequencing

ScRNA-seq libraries were prepared from viably thawed PBMCs single-cell samples prepared in the previous section according to the manufacturer’s (10x Genomics) instructions. CellPlex kit (10x Genomics, 1000261), which allows the pooling of samples before GEMs generation by labeling samples with unique cell multiplexing oligos (CMOs), was used to multiplex samples. The pooled samples were used to generate GEMs, followed by RT-PCR steps and cDNA amplification using Next GEM single cell 3’v3.1 kits (10x Genomics, 1000268). Following the cDNA amplification step, size selection beads were used to separate the CMO and gene expression (GEX) cDNAs, that were then used to prepare the CMO and GEX libraries, respectively. The final CMO and GEX libraries were then pooled and sequenced according to 10x Genomics sequencing parameters using Novaseq S4 PE100 (Illumina) kits for comprehensive transcriptome profiling.

### Single cell profiling analysis

The raw FASTQ files from each multiplex sample were aligned using 10x Genomics Cell Ranger v6.1.2 ^71^ to align against a reference human genome (GRCh38) for generating raw cell-gene count matrices. The count and CMO matrices from the samples were analyzed with R (v 4.2.2) using Seurat (v 4.0.4) and other Bioconductor packages ^72, 73^. Low-quality cells were filtered using Seurat to keep only cells with >200 unique genes, >600 UMI reads, and < 20% mitochondrial UMIs. Potential doublets were marked using the doubletFinder algorithm ^74^ that identifies doublets based on neighborhood search on principal component analysis (PCA). Assuming 3.5% of doublet formation from the 10x multiplexing experiment, we performed analysis with the top 10 principal components with a neighborhood size of 0.1(pK) to predict doublets. The log-normalized cell count (scaling factor 10000) was used for selecting the top 2,000 variable genes for principal component analysis (PCA) to identify the principal components capturing the most variance across the samples. Similar cells were clustered together via Louvain clustering on the top principal components using the Seurat package, which was visualized using Uniform Manifold Approximation and Projection (UMAP) to determine the overall relationship among the cells ^75^. The cell clusters were manually annotated based on canonical cell markers described in our previous studies. The cell markers for the different cell clusters were identified by comparing target cell types with others captured in the assay using the Wilcoxon Rank Sum test (adjusted P <.10, average log2FC > 0.25, and percent cell expression > 25%).

### Differential abundance testing on single-cell neighborhoods

Differential abundance testing on single-cell data using k-nearest neighbor graphs to detect changes in composition between Long and recovered COVID single-cell neighborhoods was performed using MiloR^76^. The KNN graph was calculated from the PCA dimensions, followed by defining the neighborhood of a cell. Differential abundance in neighborhoods was tested using the negative binomial generalized linear model implementation with an adaptation of Spatial FDR correction. with FDR < 0.10.

### Regulatory network analysis

The gene regulatory networks (GRNs) for selected clusters within our dataset were estimated using pySCENIC, an implementation of SCENIC (Single-Cell Regulatory Network Inference and Clustering) ^77^. The count matrix served as the input to run GRNBoost2 and generate co-expression modules. GRNs were further inferred using the hg38_refseq-r80 (mc_v10_clust) motif database, hgnc motif annotation (v10) and pySCENIC’s default settings. Due to the stochastic nature of the GRNBoost2algorithm ^78^, slightly varying regulons are detected in each run. Hence, high-confidence regulons were filtered out if they were present in >80% of runs, while their target genes were considered if they were detected in >90% of runs. Using AUCell from pySCENIC, each cell was assigned a gene signature score (AUC) indicating the degree of transcription factor activity ^77^.

### Gene set enrichment analysis

Single sample Gene Set Enrichment analysis (ssGSEA) score represents the degree to which the genes in a particular gene set are coordinately up- or down-regulated within a cell cluster, which could serve as an indicator of gene set activity ^79^. To interrogate the pathways/gene sets that are differentially enriched in one cell type vs. other, we performed statistical analysis using the student’s t-test. The gene set enrichment analysis was performed on pathways, and gene sets were obtained from the Molecular Signatures Database ^79^. SsGSEA was performed using the escape package from the Bioconductor library, and pathways/gene sets with P < 0.01 were considered statistically significant ^80^. The exhaustion signature for NK cells was derived from the study conducted by Bi et al ^81^.

### Cell cycle scoring analysis

Cell cycle scores were calculated using the CellCycleScoring() method in Seurat, which uses a collection of genes from a study by Tirosh et al. to assign cells to either the G1 phase, S phase, or G2/M phase of the cell cycle ^82^.

### Non-Negative Matrix Factorization

Single-cell RNA-seq data (cells × genes) was decomposed into a small set of interpretable gene programs using non-negative matrix factorization implemented in the geneNMF package ^83^.

#### Viral Repertoire analysis

Reads that were not aligned to *Homo sapiens* were processed for taxonomic profiles. Taxonomic profiling was performed using Kraken2 (v.2.1.2) and mg2sc metagenomic analysis pipeline (https://github.com/julianeweller/mg2sc) using the standard database, followed by relative abundance estimation ^84^. Feature selection was performed to identify the 50 most variable genera, which were retained for subsequent downstream processing. Genera were ranked by significance using the Wilcoxon rank-sum test; the five most differentially abundant taxa per sample were then selected for further analysis.

### Cell Communication Analysis

Cellular communication analysis was performed using the CellChat platform ^85^. Cells from the recovered and Long COVID groups were isolated, and ligand-receptor (L-R) analysis was performed on the groups independently using the standard CellChat analysis. Differentially expressed signaling genes were identified using the Wilcoxon rank sum test (P< 0.05), which was followed by communication probability/strength calculation between any interacting cell types. The cell-cell communications were filtered out if they were present in a cell type/subtype with fewer than 10 cells. The number of interactions and their strengths were aggregated for each group. To compare the overall signaling structure between cells in recovered and Long COVID samples, interaction weights were used, which sum the information flow of all L-R interactions between two groups of cell types of lymphoid and myeloid lineages.

## Acknowledgments

Sequencing was carried out at the NPRC genomics core at Emory University.

## Author contributions

Conceptualization: SS, SJ, TAW, MB

Methodology: MoB

Investigation: SS, SJ, TAW, MB

Visualization: SS, SJ

Funding acquisition: TAW, MB

Project administration: JE, TAW, MoB, MB

Formal Analysis: SS, SJ, CY, SM

Supervision: MB, TAW

Writing – original draft: SS, SJ, DO, MoB, TAW, MB

Writing – review & editing: SS, SD, DO, MoB, TAW, MB, LN

## Data and materials availability

All data, code, and materials for the paper would be provided on request.

## Supplementary Materials

**Figs. S1 to S3**

**Fig S1.** Transcriptomic characterization of B cell subtypes in recovered and Long COVID cohort.

**Fig S2.** Mapping monocyte diversity and regulatory signatures in Long COVID.

**Fig S3.** Transcriptional and regulatory features of T/NK cell subsets in COVID-19.

**Table S1.** Clinical and neurocognitive profile of participants.

