## Supplementary Materials for "Single-cell profiling of innate and adaptive immune dysregulation in Long COVID"

**The PDF file includes:**

Figs. S1 to S3
Tables S1

**Supplementary Materials**

**Figs. S1 to S3**
**Fig S1.** Transcriptomic characterization of B cell subtypes in recovered and Long COVID cohort.
**Fig S2.** Mapping monocyte diversity and regulatory signatures in Long COVID.
**Fig S3.** Transcriptional and regulatory features of T/NK cell subsets in COVID-19.

**Table S1.** Clinical and neurocognitive profile of participants.

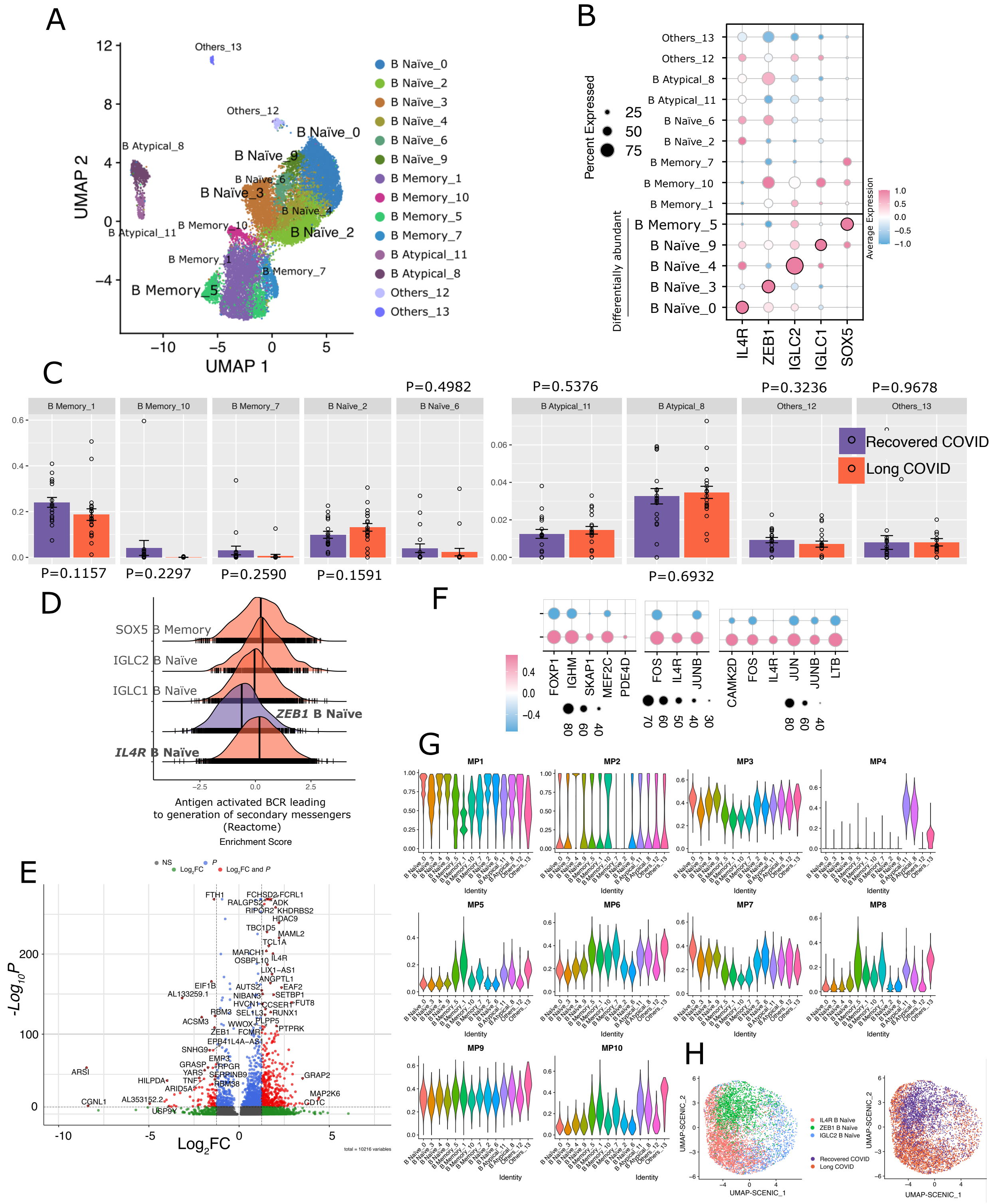
A
UMAP 2
UMAP 1
Others_13
Others_12
B Atypical_8
B Atypical_11
B Naïve_0
B Naïve_9
B Naïve_6
B Naïve_3
B Naïve_4
B Naïve_2
B Memory_10
B Memory_1
B Memory_7
B Memory_5
B Naïve_0
B Naïve_2
B Naïve_3
B Naïve_4
B Naïve_6
B Naïve_9
B Memory_1
B Memory_10
B Memory_5
B Memory_7
B Atypical_11
B Atypical_8
Others_12
Others_13
B
Percent Expressed
25
50
75
Average Expression
Differentially abundant
IL4R
ZEB1
IGLC2
IGLC1
SOX5
C
P=0.4982
P=0.5376
P=0.3236
P=0.9678
Recovered COVID
Long COVID
P=0.1157
P=0.2297
P=0.2590
P=0.1591
P=0.6932
D
SOX5 B Memory
IGLC2 B Naïve
IGLC1 B Naïve
ZEB1 B Naïve
IL4R B Naïve
Antigen activated BCR leading
to generation of secondary messengers
(Reactome)
Enrichment Score
E
-Log10P
Log2FC
F
FOXP1
IGHM
SKAP1
MEF2C
PDE4D
FOS
IL4R
JUNB
CAMK2D
FOS
IL4R
JUN
JUNB
LTB
G
MP1
MP2
MP3
MP4
MP5
MP6
MP7
MP8
MP9
MP10
Identity
H
IL4R B Naïve
ZEB1 B Naïve
IGLC2 B Naïve
Recovered COVID
Long COVID
UMAP-SCENIC_1
UMAP-SCENIC_2

**Fig. S1.**

**Transcriptomic characterization of B cell subtypes in recovered and long COVID cohort.**
**A.** UMAP projection of B cell clusters identified from single-cell RNA sequencing, showing distinct populations including B Naïve, B Memory, B Atypical, and Others. **B.** Dotplot of average gene expression for selected markers (IL4R, ZEB1, IGLC1, IGLC2, SOX5) across B cell subtypes. **C.** Comparative analysis of proportion of B-cell subtypes between long and recovered COVID groups. Bars indicate the mean proportion of each subtype normalized to the total B cell compartment per patient. Purple bars represent recovered COVID; orange bars represent long COVID group. Statistical significance assessed using Wilcoxon rank-sum test. **D.** Enrichment analysis showing Reactome pathway "Antigen activated BCR leading to generation of secondary messengers" enriched in specific B cell subsets. **E.** Volcano plot depicting differentially expressed genes comparing IL4R+ against ZEB1+ naïve B cell, highlighting genes with significant fold changes and p-values. Positive log fold change (LFC) values highlight genes enriched in IL4R+ naïve B cells, while negative LFC values indicate enrichment in ZEB1+ naïve B cells. **F.** Violin plots showing AUCell scores for gene modules derived from non-negative matrix factorization (NMF). Each violin illustrates the distribution of module activity across individual cells. **G.** Dot plot of genes overlapping between NMF modules 3 and 7 and selected MSigDB pathways, highlighting shared transcriptional programs and functional enrichment. **H.** UMAP projection based on regulon activity, where each regulon is treated as a feature to compute reduced dimensions, revealing the regulatory landscape of B cell subtypes. Left: grouped by selected B cell subtypes. Right: grouped by disease cohort (recovered vs long COVID).

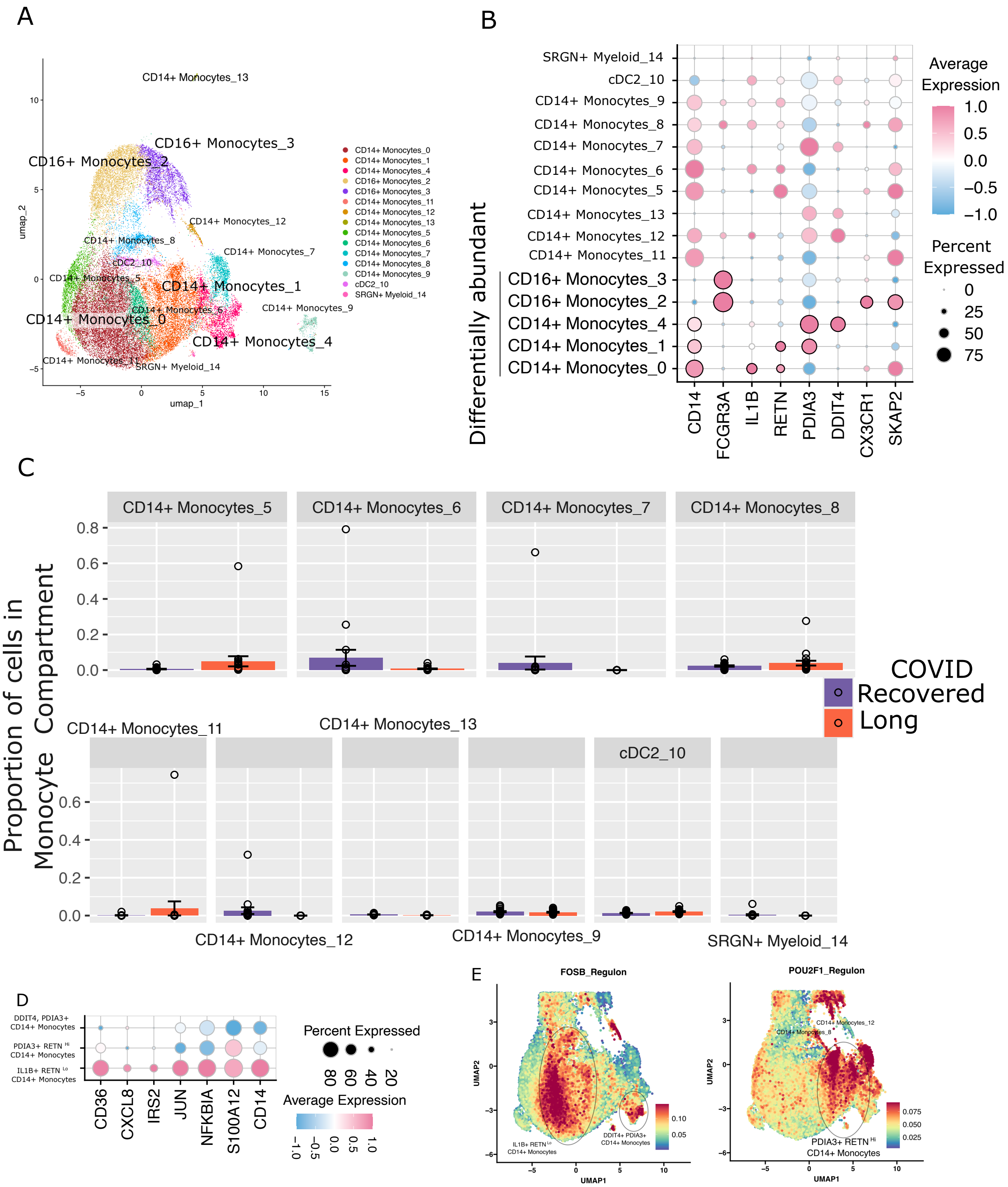
A
B
C
D
E
Differentially abundant
Average Expression
Percent Expressed
Proportion of cells in Monocyte Compartment
COVID
Recovered
Long
FOSB_Regulon
POU2F1_Regulon

**Fig. S2.**

**Mapping monocyte diversity and regulatory signatures in long COVID. A.** UMAP projection of monocyte and dendritic cell clusters, including CD14$^+$ Monocytes, CD16$^+$ Monocytes, cDC2, and SRGN$^+$ Myeloid cells. **B.** Dotplot of average gene expression for selected markers (*CD14, FCGR3A, IL1B, RETN, PDIA3, DDIT4, CX3CR1, SKAP2*) across myeloid cell subtypes. **C.** Comparative analysis of proportion of myeloid cell subtypes between long and recovered COVID groups. Bars indicate the mean proportion of each subtype normalized to the total myeloid cell compartment per patient. Purple bars represent recovered COVID; orange bars represent long COVID group. Statistical significance assessed using Wilcoxon rank-sum test. **D.** Dot plot of cytokine signaling-related genes (e.g. *CD36, CXCL8, IRS2, JUN, NFKBIA, S100A12, CD14*), showing percent and average expression across monocyte subsets. **E.** UMAP projections based on gene expression, highlighting AUCell enrichment scores for two transcriptional regulons – FOSB and POU2F1.

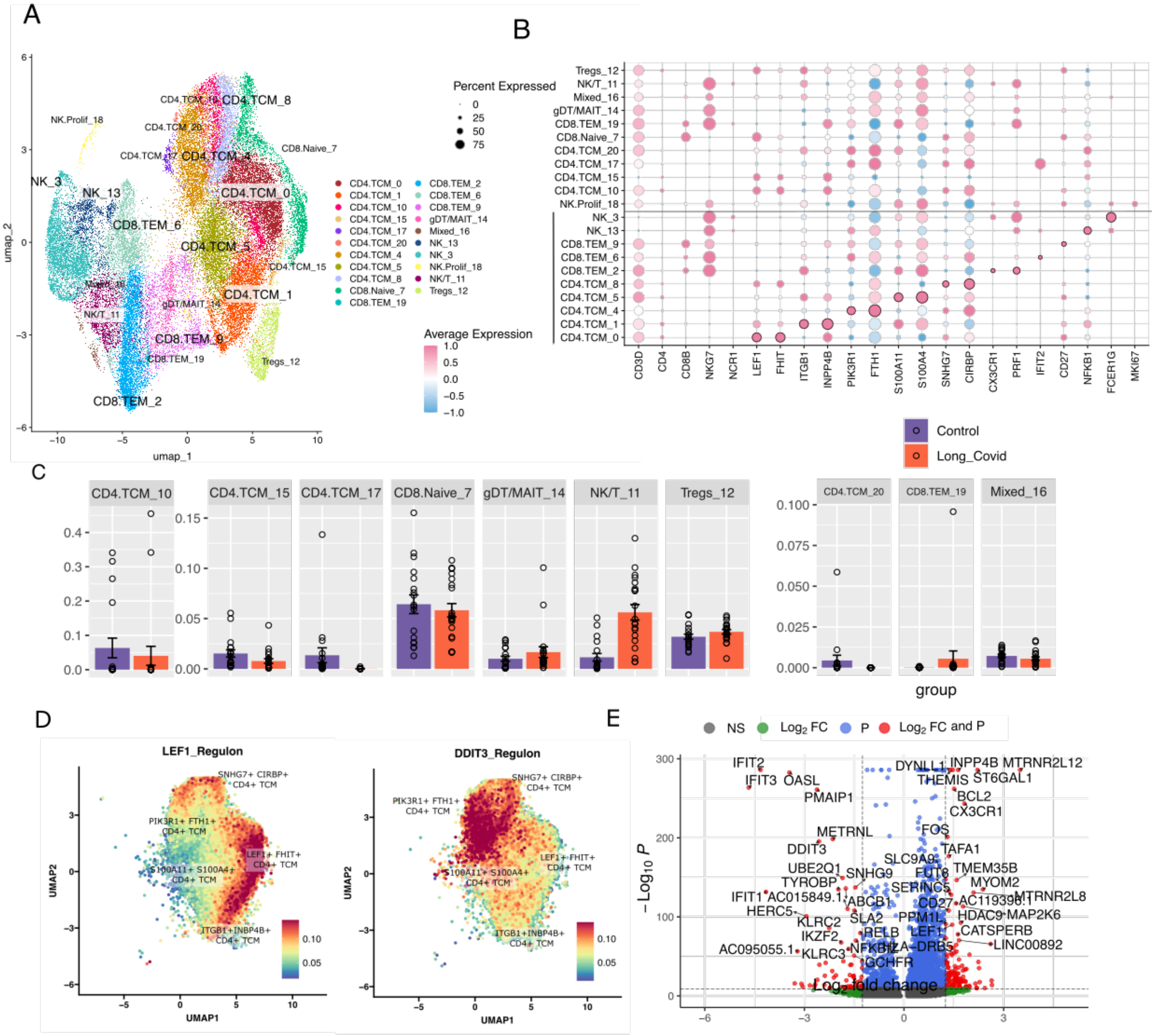

A
B
C
D
E
umap_1
umap_2
Percent Expressed
0
25
50
75
Average Expression
1.0
0.5
0.0
−0.5
−1.0
CD4.TCM_0
CD4.TCM_1
CD4.TCM_10
CD4.TCM_15
CD4.TCM_17
CD4.TCM_20
CD4.TCM_4
CD4.TCM_5
CD4.TCM_8
CD8.Naive_7
CD8.TEM_19
CD8.TEM_2
CD8.TEM_6
CD8.TEM_9
gDT/MAIT_14
Mixed_16
NK_13
NK_3
NK.Prolif_18
NK/T_11
Tregs_12
CD3D
CD4
CD8B
NKG7
NCR1
LEF1
FHIT
ITGB1
INPP4B
PIK3R1
FTH1
S100A11
S100A4
SNHG7
CIRBP
CX3CR1
PRF1
IFIT2
CD27
NFKB1
FCER1G
MKI67
Control
Long_Covid
group
LEF1_Regulon
DDIT3_Regulon
UMAP1
UMAP2
SNHG7+ CIRBP+ CD4+ TCM
PIK3R1+ FTH1+ CD4+ TCM
LEF1+ FHIT+ CD4+ TCM
S100A11+ S100A4+ CD4+ TCM
ITGB1+INBP4B+ CD4+ TCM
NS
Log₂ FC
P
Log₂ FC and P
− Log₁₀ P
Log₂ fold change
IFIT2
IFIT3
OASL
PMAIP1
METRNL
DDIT3
UBE2Q1
TYROBP
SNHG9
IFIT1
AC015849.1
ABCB1
HERC5
KLRC2
SLA2
IKZF2
RELB
AC095055.1
KLRC3
NFKBIZ
GCHFR
DYNLL1
INPP4B
MTRNR2L12
THEMIS
ST6GAL1
BCL2
CX3CR1
FOS
TAFA1
SLC9A9
FUT8
TMEM35B
SERINC5
MYOM2
CD27
AC119396.1
MTRNR2L8
PPM1L
HDAC9
MAP2K6
LEF1
CATSPERB
HLA−DRB5
LINC00892

**Fig. S3.**

**Transcriptional and regulatory features of T/NK cell subsets in COVID-19. A.** UMAP projection of T cell clusters including $CD4^+$ TCM, $CD8^+$ TEM, $CD8^+$ Naive, Tregs, NK, NK/T, NK.Prolif, γδ/MAIT, and mixed populations. **B.** Dot plot showing average expression and percent expression of canonical T cell markers and signaling genes (e.g., *CD3D, CD4, CD8B, NKG7, LEF1, CX3CR1, PRF1, IFIT2, CD27, NFKB1, MKI67*) across T cell subsets. **C.** Comparative analysis of proportion of T/NK cell subtypes between long and recovered COVID groups. Bars indicate the mean proportion of each subtype normalized to the total T/NK lymphoid cell compartment per patient. Purple bars represent recovered COVID; orange bars represent long COVID group. Statistical significance assessed using Wilcoxon rank-sum test. **D.** UMAP projections based on gene expression, highlighting AUCell enrichment scores for two transcriptional regulons – LEF1 and DDIT3. **E.** Volcano plot of differentially expressed genes comparing $CD8^+$ TEM cells between long COVID and recovered cohorts. Positive log fold change (LFC) values indicate genes enriched in long COVID, while negative LFC values indicate enrichment in recovered individuals.

**Table S1.**

| | Recovered COVID | Long COVID | P |
|---|---|---|---|
| | (N=18) | (N=20) | |
| **PROMIS® v2.0 Cognitive Function Test** | | | |
| Mean (SD) | 34.9 (6.13) | 25.2 (9.28) | <0.001 |
| Median [Min, Max] | 38.0 [24.0, 40.0] | 24.5 [9.00, 40.0] | |
| **Modified fatigue impact scale, MFIS** | | | |
| Mean (SD) | 21.3 (22.7) | 37.9 (21.9) | 0.0288 |
| Median [Min, Max] | 16.0 [0, 75.0] | 37.0 [0, 82.0] | |
| **Global function deficit severity (based on z-score)** | | | |
| none | 3 (16.7%) | 1 (5.0%) | 0.429 |
| mild | 7 (38.9%) | 7 (35.0%) | |
| moderate | 1 (5.6%) | 4 (20.0%) | |
| severe | 7 (38.9%) | 8 (40.0%) | |
| **Memory deficit severity (based on z-score)** | | | |
| none | 0 (0%) | 0 (0%) | 0.0534 |
| mild | 10 (55.6%) | 4 (20.0%) | |
| moderate | 4 (22.2%) | 5 (25.0%) | |
| severe | 4 (22.2%) | 11 (55.0%) | |
| **Attention deficit severity (based on z-score)** | | | |
| none | 4 (22.2%) | 0 (0%) | 0.127 |
| mild | 4 (22.2%) | 4 (20.0%) | |
| moderate | 3 (16.7%) | 3 (15.0%) | |
| severe | 7 (38.9%) | 13 (65.0%) | |
| **Executive function deficit severity (based on z-score)** | | | |
| none | 5 (27.8%) | 5 (25.0%) | 0.816 |
| mild | 11 (61.1%) | 12 (60.0%) | |
| moderate | 2 (11.1%) | 2 (10.0%) | |
| severe | 0 (0%) | 1 (5.0%) | |
| **Generalized anxiety disorder scale (GAD-7)** | | | |
| Mean (SD) | 9.83 (9.04) | 16.3 (14.9) | 0.115 |
| Median [Min, Max] | 8.00 [0, 28.0] | 11.5 [0, 44.0] | |
| **Post-traumatic stress disorder checklist (PCL-C),** | | | |
| Mean (SD) | 34.6 (14.1) | 37.9 (17.3) | 0.522 |
| Median [Min, Max] | 29.5 [17.0, 62.0] | 36.0 [17.0, 83.0] | |
| **CES Depression Scale** | | | |
| Mean (SD) | 18.8 (13.0) | 20.6 (11.3) | 0.658 |
| Median [Min, Max] | 15.0 [0, 41.0] | 17.0 [4.00, 44.0] | |
| **Hopkins Verbal Learning Test immediate (HVLT-I, z-score)** | | | |

| | | | |
|---|---|---|---|
| Mean (SD) | -0.172 (1.12) | -1.24 (0.691) | 0.00161 |
| Median [Min, Max] | -0.370 [-1.77, 1.77] | -1.35 [-2.31, 0.24] | |
| **Hopkins Verbal Learning Test immediate delayed (HVLT-D, z-score)** | | | |
| Mean (SD) | 0.0694 (1.12) | -1.02 (0.851) | 0.00215 |
| Median [Min, Max] | 0.192 [-1.79, 1.96] | -1.02 [-2.36, 0.714] | |
| **Craft Story immediate recall (CRAFTVRS, z-score)** | | | |
| Mean (SD) | -1.21 (0.897) | -1.32 (1.33) | 0.765 |
| Median [Min, Max] | -1.15 [-3.25, 0.156] | -1.72 [-3.00, 1.45] | |
| **Craft story delayed recall (CRAFDVRS, z-score)** | | | |
| Mean (SD) | -1.24 (0.802) | -1.40 (0.987) | 0.593 |
| Median [Min, Max] | -1.25 [-2.70, -0.00800] | -1.59 [-2.61, 1.02] | |
| **Montreal Cognitive Assessment attention (MoCA ATTN, zscore)** | | | |
| Mean (SD) | -2.14 (2.18) | -3.83 (1.92) | 0.016 |
| Median [Min, Max] | -1.44 [-7.32, 0.0290] | -4.01 [-7.32, -0.705] | |
| **Montreal Cognitive Assessment (MoCA)** | | | |
| Mean (SD) | -1.55 (1.77) | -1.84 (1.22) | 0.563 |
| Median [Min, Max] | -1.10 [-4.64, 1.43] | -1.65 [-4.74, -0.206] | |
| **Number span forward (DIGFORCT, z-score)** | | | |
| Mean (SD) | 0.431 (1.33) | -0.131 (0.977) | 0.151 |
| Median [Min, Max] | 0.440 [-1.47, 2.65] | -0.267 [-1.47, 2.90] | |
| **Number span backward (DIGBACCT, z-score)** | | | |
| Mean (SD) | 0.216 (1.78) | -0.152 (1.13) | 0.459 |
| Median [Min, Max] | -0.286 [-1.78, 3.33] | -0.393 [-1.73, 3.22] | |
| **Oral Trail Making Part B (OT-B, Zscore)** | | | |
| Mean (SD) | 0.465 (0.582) | 1.35 (0.619) | <0.001 |
| Median [Min, Max] | 0.444 [-0.579, 1.52] | 1.39 [-0.406, 2.07] | |
| **Cognitive Mental Control** | | | |
| Mean (SD) | 25.2 (6.85) | 17.8 (3.76) | <0.001 |
| Median [Min, Max] | 24.5 [13.0, 38.0] | 18.5 [10.0, 24.0] | |
| **Any of the following symptoms that were new or worsened since COVID-19 infection(s)** | | | |

| | | | |
|---|---|---|---|
| **Dyspnea (shortness of breath or trouble breathing)** | | | |
| No | 14 (77.8%) | 7 (35.0%) | 0.0203 |
| Yes | 4 (22.2%) | 13 (65.0%) | |
| **Fatigue** | | | |
| No | 15 (83.3%) | 5 (25.0%) | 0.00107 |
| Yes | 3 (16.7%) | 15 (75.0%) | |
| **PEM, Post-exertional malaise (delayed fatigue after activities or exercise)** | | | |
| No | 17 (94.4%) | 13 (65.0%) | 0.0681 |
| Yes | 1 (5.6%) | 7 (35.0%) | |
| **Dizziness** | | | |
| No | 18 (100%) | 15 (75.0%) | 0.0725 |
| Yes | 0 (0%) | 5 (25.0%) | |
| **Headache** | | | |
| No | 16 (88.9%) | 12 (60.0%) | 0.0989 |
| Yes | 2 (11.1%) | 8 (40.0%) | |
| **Palpitations** | | | |
| No | 18 (100%) | 17 (85.0%) | 0.267 |
| Yes | 0 (0%) | 3 (15.0%) | |
| **Trouble concentrating** | | | |
| No | 18 (100%) | 3 (15.0%) | <0.001 |
| Yes | 0 (0%) | 17 (85.0%) | |
| **Language, Word finding difficulties** | | | |
| No | 16 (88.9%) | 6 (30.0%) | <0.001 |
| Yes | 2 (11.1%) | 14 (70.0%) | |
| **Memory loss** | | | |
| No | 17 (94.4%) | 4 (20.0%) | <0.001 |
| Yes | 1 (5.6%) | 16 (80.0%) | |
| **Difficulties multitasking** | | | |
| No | 18 (100%) | 10 (50.0%) | 0.00177 |
| Yes | 0 (0%) | 10 (50.0%) | |
| **Trouble staying asleep** | | | |
| No | 16 (88.9%) | 9 (45.0%) | 0.0122 |
| Yes | 2 (11.1%) | 11 (55.0%) | |
| **Unrefreshing sleep** | | | |
| No | 16 (88.9%) | 10 (50.0%) | 0.026 |
| Yes | 2 (11.1%) | 10 (50.0%) | |
| **Hair loss** | | | |

| | | | |
|---|---|---|---|
| No | 18 (100%) | 14 (70.0%) | 0.0369 |
| Yes | 0 (0%) | 6 (30.0%) | |

**Table S1. Clinical and neurocognitive profile of participants.** Data include standardized neurocognitive test scores, symptom severity measures, and the patient-reported prevalence of Long COVID symptoms (dichotomous).
